# Structure of the SHOC2–MRAS–PP1C complex provides insights into RAF activation and Noonan syndrome

**DOI:** 10.1101/2022.05.10.491335

**Authors:** Daniel A. Bonsor, Patrick Alexander, Kelly Snead, Nicole Hartig, Matthew Drew, Simon Messing, Lorenzo I. Finci, Dwight V. Nissley, Frank McCormick, Dominic Esposito, Pablo Rodriguez-Viciana, Andrew G. Stephen, Dhirendra K. Simanshu

## Abstract

SHOC2 acts as a strong synthetic lethal interactor with MEK inhibitors in multiple KRAS cancer cell lines. SHOC2 forms a heterotrimeric complex with MRAS and PP1C that is essential for regulating RAF and MAPK-pathway activation by dephosphorylating a specific phosphoserine on RAF kinases. Here we present the high-resolution crystal structure of SHOC2-MRAS-PP1C (SMP) complex and apo-SHOC2. Our structures reveal that SHOC2, MRAS and PP1C form a stable ternary complex where all three proteins synergistically interact with each other. Our results show that dephosphorylation of RAF substrates by PP1C is enhanced upon interacting with SHOC2 and MRAS. The SMP complex only forms when MRAS is in an active state and is dependent on SHOC2 functioning as a scaffolding protein in the complex by bringing PP1C and MRAS together. Our results provide structural insights into the role of the SMP complex in RAF activation, how mutations found in Noonan syndrome enhance the complex formation and reveal new avenues for therapeutic interventions.

## INTRODUCTION

The mitogen-activated protein kinases (MAPK) signaling pathway comprises the RAF, MEK, and ERK protein kinases, constituting a critical effector cascade used by the RAS proteins to regulate cell growth, survival, proliferation, and differentiation^1^. Aberrant activation of the MAPK signaling pathway is one of the most common drivers in human cancer and is responsible for multiple developmental disorders known as RASopathies^2,3^. Within this signaling pathway, the regulation of RAF kinases is a complex process that involves protein and lipid interactions, subcellular localization, and multiple phosphorylation/dephosphorylation events^4^. RAF kinases are held in an autoinhibited state by the 14-3-3 family of phosphoserine/phosphothreonine-binding proteins which bind to RAF using two phosphorylation-dependent 14-3-3 binding sites^5,6^. In RAF kinase these two phosphorylation sites are located within Conserved Region 2 (CR2) at the N-terminal end of the kinase domain (ARAF S214, BRAF S365, CRAF/RAF1 S259, hereby referred to as CR2-pS), and in Conserved Region 3 (CR3) in the C-terminal tail that follows the kinase domain (ARAF S582, BRAF S729, CRAF S621, hereby referred to CR3-pS). RAF Kinase activation requires active RAS binding to the RAS-binding domain (RBD) and membrane-anchoring cysteine-rich domain (CRD) of RAF^7,8^. Dephosphorylation of CR2-pS is a critical step in the RAF activation process as it prevents 14-3-3 binding at this site. This step allows the released kinase domain to dimerize, forming an active dimeric RAF complex that is stabilized by a 14-3-3 dimer bound to the CR3-pS sites of each RAF kinase. CRAF/RAF1 mutations (S257L and P261S) around the CR2-pS259 14-3-3 binding site are frequently detected in RASopathy Noonan syndrome (NS)^9^. These mutations have been suggested to enhance CRAF activation by disrupting 14-3-3 binding to the S259 site, underscoring the critical role of this step in RAF and MAPK-pathway regulation.

The dephosphorylation of CR2-pS is mediated by a heterotrimeric complex comprised of SHOC2, MRAS, and PP1C (SMP complex), where each of the three proteins plays indispensable roles in the proper function of the complex^10,11^. SHOC2 was initially identified in *C. elegans* as a positive modulator of the MAPK pathway^12,13^. It is a ubiquitously expressed protein composed primarily of predicted leucine-rich repeats (LRRs). N-terminal to the LRR domains, SHOC2 contains a ∼90-residue long sequence that is predicted to be intrinsically disordered and has been suggested to be necessary for complex formation with MRAS and PP1C^11,14^. Germline mutations in SHOC2 (S2G, M173I, and Q269H/H270Y) have been detected in NS^11,15–17^. SHOC2 plays a vital role in transformation, metastasis, epithelial-to-mesenchymal transition, and MAPK pathway inhibitor resistance^18–21^. Multiple genome-scale, single-gene CRISPR/Cas9 fitness screens in human cancer cells have suggested selective dependency of RAS mutant cells on SHOC2^20,22–24^. SHOC2 was also identified as the strongest synthetic lethal target in the presence of MEK inhibitors in KRAS mutant lung and pancreatic cancer cell lines^19^. Thus, SHOC2 may provide a unique therapeutic opportunity within the RTK-RAS-MAPK pathway in oncogenic RAS cells.

The SMP complex formation is initiated following MRAS activation as SHOC2 and PP1C bind only to MRAS-GTP^25^. The canonical RAS family members HRAS, KRAS, and NRAS, also bind SHOC2, although with considerably lower affinity than MRAS^26^. The nature of the selectivity for MRAS is not known. MRAS shares ∼50% sequence identity with the canonical RAS proteins and contains an extra ten amino acids at the N-terminus. Activating mutations in MRAS are very rare in cancer; however, gain-of-function mutations (G23V, T68I, Q71R) in MRAS have been identified in NS patients^27,28^. In the SMP complex, PP1C provides the enzymatic activity for dephosphorylation. PP1C is a class of serine/threonine phosphatases with three highly conserved isoforms (PP1CA, PP1CB, and PP1CC with >90% sequence identity) that are ubiquitously expressed and catalyze the dephosphorylation of a substantial fraction of phosphoserine/threonine in all eukaryotic cells^29–31^. Mutations in the PP1CB isoform have been found in NS, and these residues are conserved in other PP1C isoforms^32–35^.

To understand how SHOC2, MRAS, and PP1C proteins assemble to form a ternary complex that regulates dephosphorylation of the RAF CR2-pS and how RASopathy mutations impact complex formation, we solved the structure of the SMP complex at 2.17 Å. Structural and mutational analysis provide a rationale for MRAS selectivity versus canonical RAS isoforms and the impact of NS mutations on the SMP complex assembly. SHOC2 and MRAS enhance the dephosphorylation activity of PP1C upon complexation towards RAF substrates. Analysis of the protein-protein interfaces in the SMP complex and mutagenesis studies provide insights into complex assembly and potential sites that could be exploited using structure-based drug discovery approaches.

## RESULTS

### Assembly of the SHOC2-MRAS-PP1CA complex

The pre-assembled human SMP complex was purified by co-expressing SHOC2, MRAS (Q71L or wild-type) and PP1CA from a single plasmid (Fig. 1a) along with the chaperone SUGT1 in baculovirus-infected insect cells, as described previously^36^. Nucleotide analysis of the purified SMP complex showed that MRAS is bound to GTP (Supplementary Fig. 1a). Using surface plasmon resonance (SPR), we measured the affinity of SMP complexation with the individually purified proteins and with MRAS in GDP or GMPPNP-bound states. Low level, transient association of PP1CA was observed with SHOC2 but could not be quantitated (Fig. 1b-c). Stable SMP complexation was only observed when PP1CA and MRAS-GMPPNP were flowed over Avi-tagged SHOC2 present on the chip surface, showing a K_D_ of ∼120 nM (Fig. 1c); no complex was formed when PP1CA was combined with MRAS-GDP (Fig. 1b). We also measured the affinity of this interaction by isothermal titration calorimetry (ITC) and observed a similar affinity of ∼350 nM, despite a higher salt concentration required for the ITC experiments (Supplementary Fig. 1b). We used the MRAS-Q71L mutant for our structural work as SPR measurements using this mutant showed SMP complex formation with a ∼5-fold higher affinity (K_D_ =26 nM) than that of WT MRAS (Fig. 1d). SPR measurement using KRAS-GMPPNP, HRAS-GMPPNP or NRAS-GMPPNP in place of MRAS-GMPPNP showed ternary complex formation (SKP, SHP or SNP) with an apparent K_D_ of 0.7 µM, 2 µM, and 4 µM, respectively, due to a substantial increase in both on-rate and off-rate (Fig. 1e-g and Supplementary Fig 1c-d). Thus, a 7-40-fold higher affinity of MRAS over RAS isoforms for complex formation confirmed that MRAS is the preferred partner of SHOC2 and PP1C.

**Figure 1:**
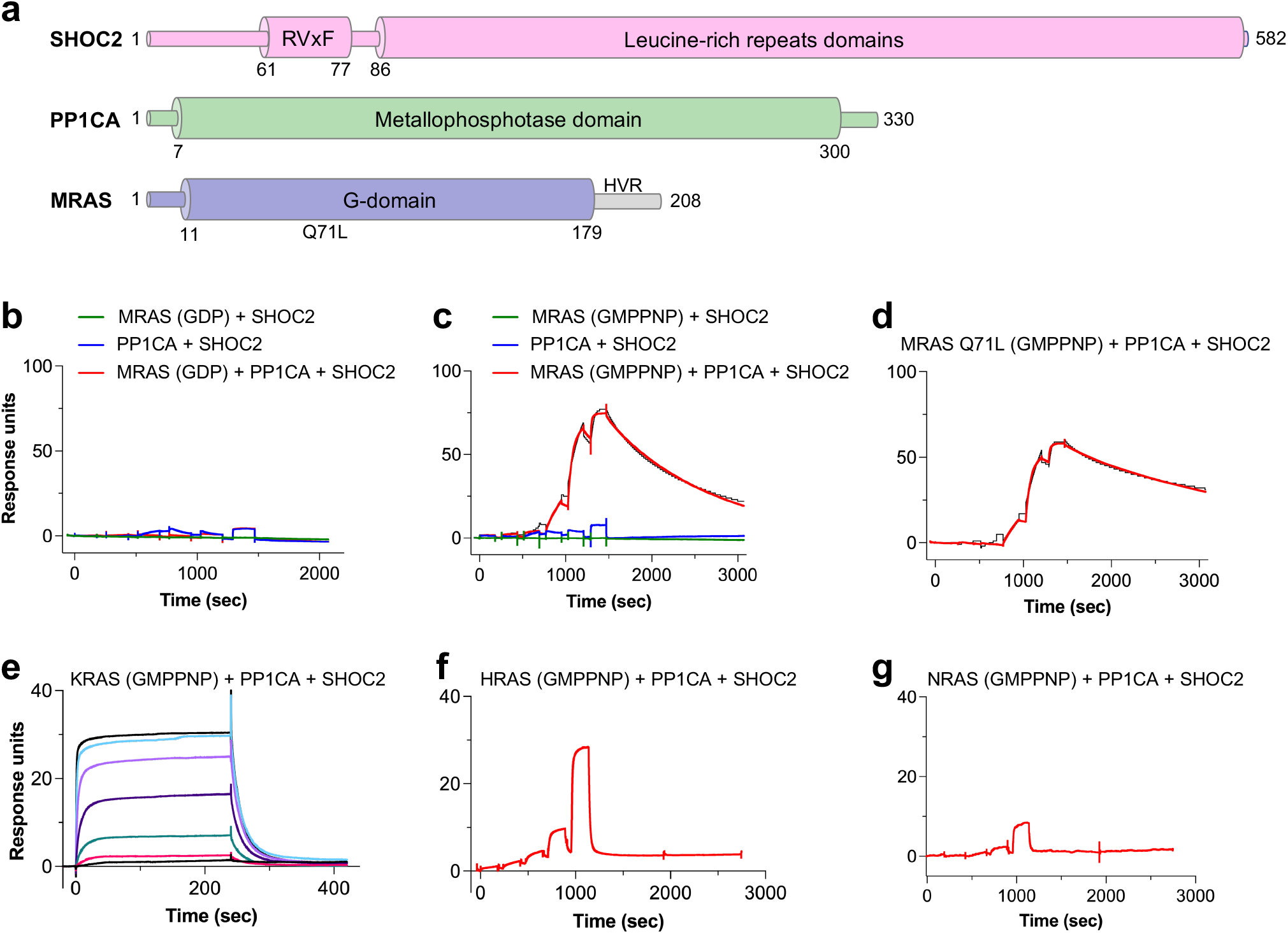
Assembly, activity, and selectivity of the SMP complex. **a** Domain architecture of SHOC2, MRAS and PP1CA. Full-length SHOC2 and PP1CA, and the G-domain of MRAS (1-179) were used for the structure determination. **b, c,** Single-cycle kinetic titration SPR binding experiments were performed on immobilized avi-tagged SHOC2 with 3-fold dilutions of 1 µM MRAS (green), PP1CA (blue), and MRAS with PP1CA (red). All experiments were either conducted with (**b)** MRAS_GDP_ or (**c)** MRAS_GMPPNP_. The data were fit to a 1:1 kinetic model (black). SMP complex assembly only occurred with MRAS bound to GMPPNP and in the presence of PP1CA. **d** Single-cycle kinetic analysis was performed on immobilized avi-tagged SHOC2 with 3-fold serial dilutions of 1 μM MRAS_Q71L-GMPPNP_ and 1 μM PP1CA (red). The data were fit to a 1:1 kinetic model (black). **e** Assembly of the SKP (SHOC2-KRAS-PP1CA) complex was measured by SPR kinetic analysis. 2-fold dilutions of 5 µM KRAS_GMPPNP_ and 5 µM PP1CA were injected over immobilized avi-tagged SHOC2. **f, g** Assembly of the SHP (SHOC2-HRAS-PP1CA) or SNP (SHOC2-NRAS-PP1CA) complexes were measured by SPR single-cycle kinetic analysis. 2-fold dilutions of **(f)** 5 µM HRAS_GMPPNP_ and 5 µM PP1CA or **(g)** 5 µM NRAS_GMPPNP_ and 5 µM PP1CA were injected over immobilized avi-tagged SHOC2.

### Structural description of the SMP complex

To understand how SHOC2, MRAS and PP1CA interact with each other, the structure of the SMP complex was determined to a resolution of 2.17 Å (Fig. 2a-b and Supplementary Table 1). In the crystal, two copies of the SMP complex are present in the asymmetric unit, labeled SMP1 and SMP2. The superposition of these two complexes showed an almost identical arrangement of three proteins (Fig. 2c and Supplementary Fig. 2a). MRAS and PP1CA are nearly identical in the two complexes (Supplementary Fig. 2a), however, the SHOC2 molecules differ between the two SMP complexes present in the asymmetric unit (Fig. 2c and Supplementary Fig. 2a-b). The C-terminus of SHOC2 in one of the SMP complexes forms additional contacts with MRAS and PP1CA (Fig. 2c and Supplementary Fig. 2a-2c). This distortion in SHOC2 is propagated and amplified causing the C-terminal helix at the end of the LRRs to move 10 Å towards MRAS (Fig. 2c and Supplementary Fig. 2a). Several SHOC2 structure predictions suggested a flexible hinge around LRR 13-15^12,14,15,18^. Our structure is consistent with this prediction that SHOC2 contains a flexible hinge, but it is within LRR 10 (residues 308-331) (Supplementary Fig. 2b-c).

**Figure 2.**
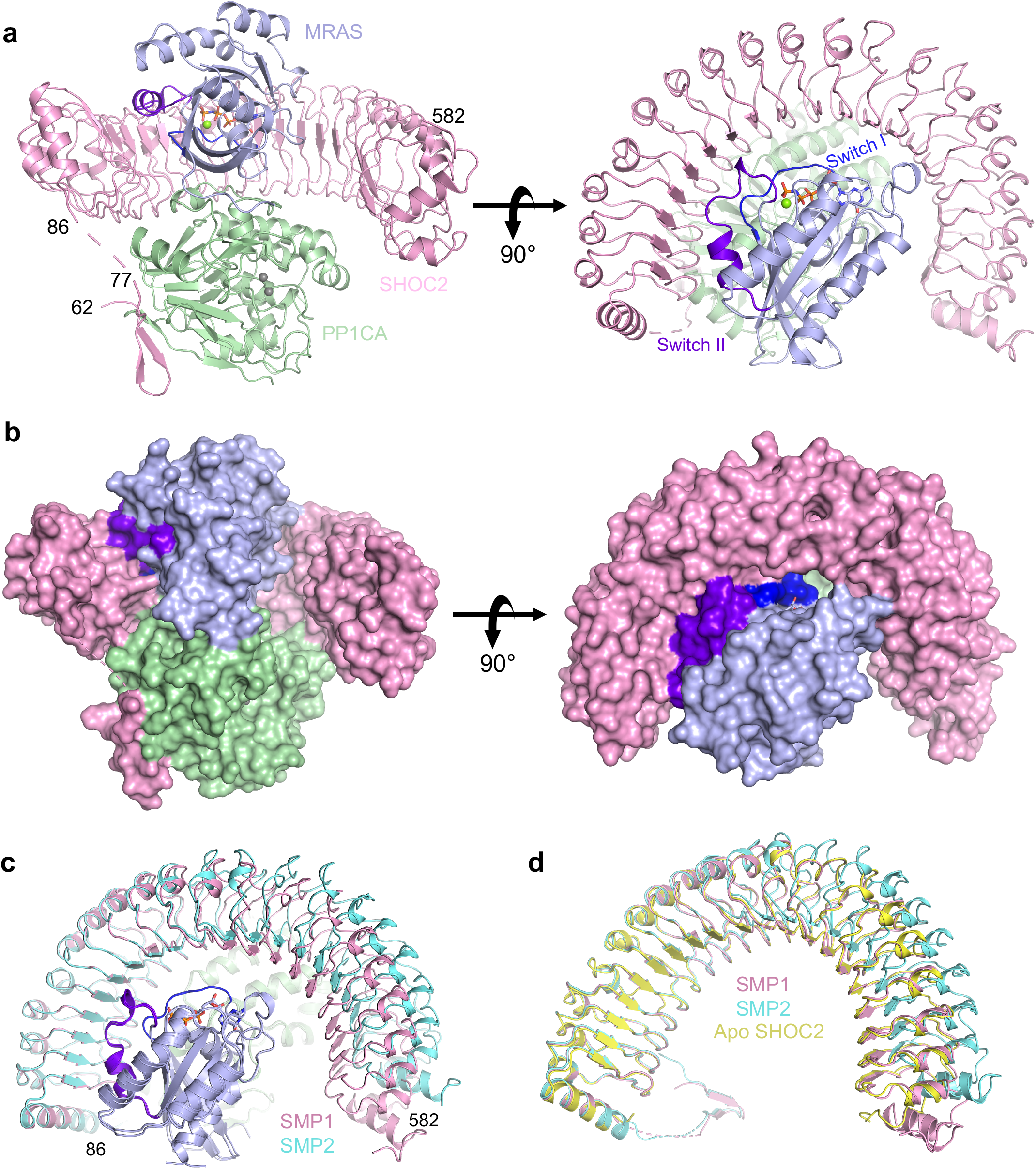
The 2.17 Å structure of the SMP complex. **a, b** The overall structure of the SMP complex is shown in **a** cartoon and **b** surface representation in two different views. SHOC2 and PP1CA are colored pink and green, respectively. MRAS is colored blue with the switch I and switch II regions highlighted in dark blue and purple, respectively. GMPPNP is shown as sticks, and Mg^2+^ (green) and Mn^2+^ (gray) ions as spheres. Active site containing Mn^2+^ ions is within the black circle. **c** Superposition of the two SMP complexes present in the asymmetric subunit in cartoon form. Both chains of MRAS and PP1CA are in the same color, while the two SHOC2 chains are colored pink and cyan. **d** Superposition of Apo-SHOC2 (yellow) onto the two SHOC2 chains from the SMP complex.

To determine if SHOC2 undergoes conformational changes upon SMP complex assembly, we solved the structure of apo-SHOC2_58-564_ at 2.4 Å (Supplementary Table 1). In the SHOC2_58-564_ structure, all LRRs and the helix that caps the N-terminal end of LRR are well defined. We do not observe any residues (58-86) prior to the N-terminal capping helix, suggesting no interaction of these residues with LRRs. Comparative structural analysis showed apo-SHOC2_58-564_ superimposes with a lower RMSD (0.57 Å) onto the SMP1 complex (Fig. 2d). SHOC2 in SMP1 forms extra contacts with MRAS and PP1CA (Fig. 2d and Supplementary Fig. 2a, 2d), which are important for binding and are described below. Our subsequent structural analysis is based on this SMP complex.

As expected, MRAS in the SMP complex adopts the highly conserved G-domain fold in an active state. Since no structure of human MRAS is available, we used the structure of mouse MRAS_GMPPNP_ (PDB ID 1X1S, 97% identity) to understand structural similarities and differences with human MRAS present in the SMP complex^37^. Superposition of mouse MRAS_GMPPNP_ with MRAS present in the SMP complex reveals significant differences in the two switch regions (Supplementary Fig. 2e). In the apo-MRAS structure, the switch I region (residues 36-50) is in the open solvent-exposed State I conformation, while this region in the SMP complex is in the closed State II conformation as observed in structures of other RAS-effector complexes^1,6,7^. Multiple residues of the switch II region (residues 69-73), which are typically disordered in the apo-MRAS structure, are ordered in the SMP complex. The Q71L mutation, which increases SMP complexation by ∼5-fold, likely aids the formation of a helical loop that contributes additional interactions from switch II to the SMP complex. Structural comparison of PP1CA in the SMP complex with human apo-PP1CA (PDB ID 4MOV) shows no major structural changes in PP1CA upon complexation with SHOC2 and MRAS (Supplementary Fig. 2f)^38^. Among the PP1C isoforms, the C-terminal tail shows high sequence diversity (residues 300-330) and is proposed to function as an inhibitor when phosphorylated at T320 (in PP1CA) and complexed with other PP1C regulators^39^. The C-terminal tail of PP1CA (residues 300-330) are disordered in our structure, though electron density exists for five residues (318-322) in one molecule of PP1CA that interact with symmetry-related SHOC2 molecules. To test whether PP1C isoform specificity exists, we used ITC to measure the affinity of SMP complexation with PP1CA and PP1CB. We observed similar affinities and thermodynamics for these two isoforms signifying no PP1C isoform specificity (Supplementary Fig. 3a-b). SHOC2 present in the SMP complex reveals structural details of this protein for the first time. The N-terminal residues of SHOC2, which have been predicted to be intrinsically disordered, are not visible in the SMP complex; however, we observe residues 60-76 of SHOC2, which fold into a β-hairpin and interacts with PP1CA (see below). The C-terminal domain of SHOC2 contains twenty LRR domains, which are capped at the termini by helices resulting in a horseshoe-shaped protein (Fig. 2a). In the SMP complex, each protein interacts with the other two proteins which results in a total of ∼6200 Å^2^ of buried surface area upon complexation.

### The SHOC2-PP1CA interface

The SHOC2-PP1CA interface of the SMP complex contributes the largest amount of buried surface area of 2800 Å^2^. SHOC2 was predicted to interact with PP1CA through SILK and RVxF motifs identified in the folded region of SHOC2 between LRR 10-11 and within LRR 12 (Supplementary Fig. 4a)^18^. These short linear motifs have always been found in disordered regions of proteins and as observed in our structure, PP1CA does not contact these residues (Supplementary Fig. 4a). However, we do identify a bona fide RVxF motif within the β-hairpin of the N-terminal intrinsically disordered domain of SHOC2 that interacts with PP1CA (Fig. 2a, 3a-b). This RVxF motif (^62^PGVAF^66^) would not be recognized by RVxF prediction algorithms^40,41^. The RVxF motif buries V64 and F66 into the hydrophobic pockets on the surface of PP1CA (Fig. 3a-b), with the β-hairpin forming 5 hydrogen bonds and burying ∼1150 Å^2^ of surface area (Fig. 3c). The binding of the SHOC2 RVxF motif, like other RVxF-containing protein, to PP1CA mimics all PP1CA-Protein_RVxF_ complexes whose structures have been solved and does not alter the structure of PP1CA (Supplementary Fig. 4b). The RVxF motif is believed to function as an anchoring motif^31^. RVxF motif binding can be regulated through phosphorylation of the variable residue if a serine or threonine is present. The presence of an alanine in that position in SHOC2 prevents its regulation directly. However, residue T71 in SHOC2, which lies on the second strand of the β-hairpin, has been shown to be phosphorylated and may play a role in the regulation of the SMP complex^42^. The SHOC2 RVxF motif is conserved across higher eukaryotes (Supplementary Fig. 4c). Mutation of valine or phenylalanine drastically weakens the formation of the SMP complex as measured by SPR, with nearly a 600-fold reduction in the apparent K_D_ (Fig. 3d and Supplementary Fig. 4d). These SHOC2 mutants retain weakened SMP complexation due to the extensive contacts through the LRRs to PP1CA involving residues preceding and/or succeeding the conserved asparagine of the Highly Conserved Segment (HCS) motif (LxxLxLx<u>x</u>N(<u>x</u>)_1-2_L) in LRRs 2-5, 8-11, and 13-18 (Fig. 3e). As such, when mapped onto the surface of SHOC2, PP1CA contacts the underside of the LRR domain (Fig. 3e). This binding interface is larger than the RVxF interaction with the burial of ∼1650 Å^2^ of surface area and the formation of nine hydrogen bonds and eight salt bridges (Fig. 3f-h). The E155 SHOC2 residue forms a hydrogen bond with R188 of PP1CA. The mutation E155A results in a ∼10-fold reduction in the apparent affinity compared to wild type (Fig. 3d and Supplementary Fig. 4d). SHOC2 does not contain a SILK motif, however, it does interact with the periphery of the SILK binding pocket on PP1CA through van der Waal interactions and a hydrogen bond between R203 of SHOC2 and E54 of PP1CA (Fig. 3f, 3h and Supplementary Fig. 4e). SHOC2 does not occupy the SILK binding pocket but rather restricts access to it. The double mutation of the SILK binding pocket in PP1CB, E53A/L54A (E53 being equivalent to E54 in PP1CA due to the presence of an extra amino acid at the N-terminus, Supplementary Fig. 5a) has previously been shown to weaken complex formation^11^. This suggests that the hydrogen bond between R203 of SHOC2 and E54 of PP1CA is important (L55 of PP1CA does not contact SHOC2). In the SMP complex, the LRR domain of SHOC2 acts as a tiara interacting with the crown of PP1CA (Fig. 2a, 3e), with SHOC2 residues >20 Å away from the PP1CA active site (Fig. 2a).

**Figure 3.**
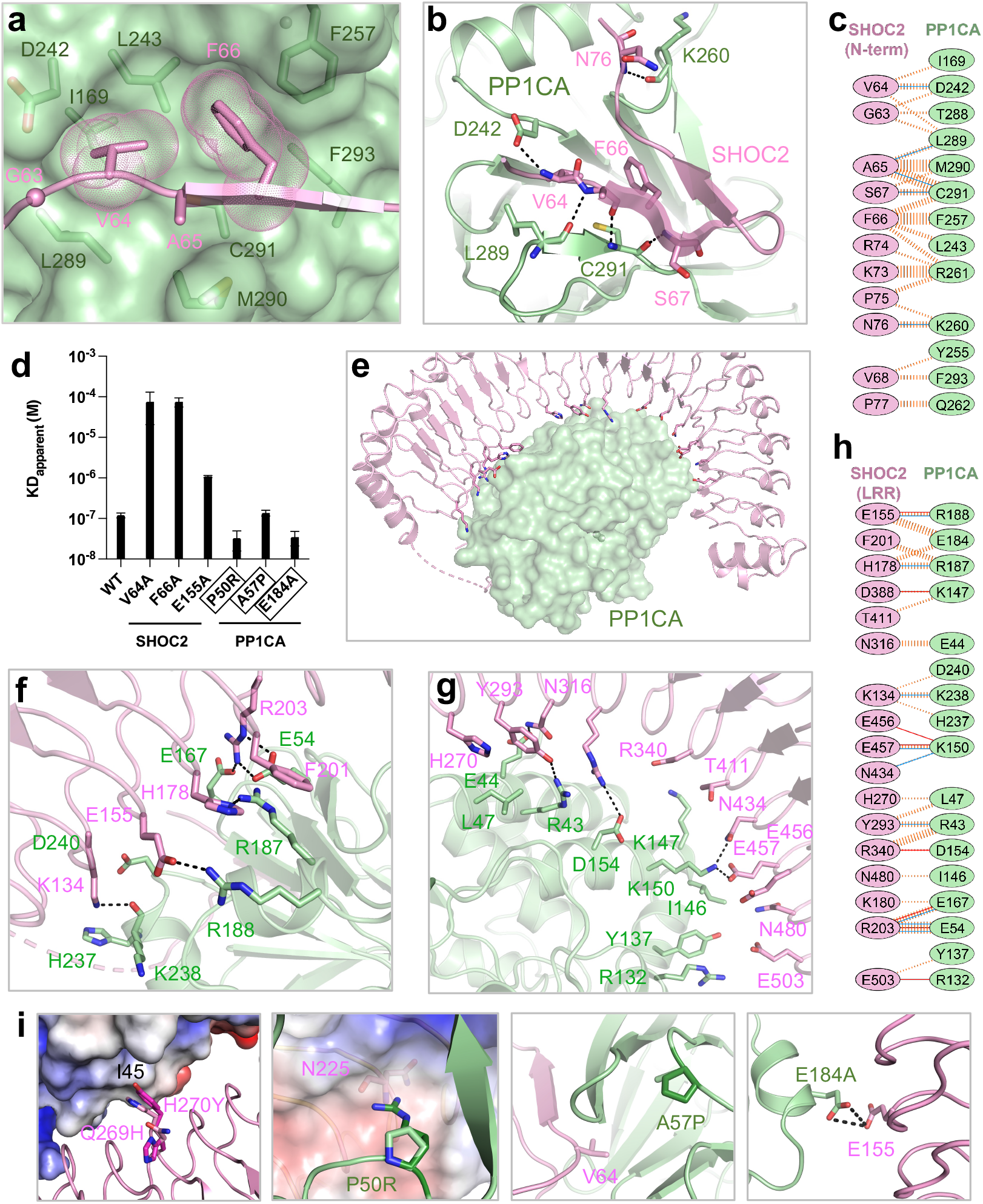
Structural and mutational analysis of the SHOC2-PP1CA interface. **a** The RVxF motif of SHOC2 (pink) bound to the surface of the RVxF binding pocket of PP1CA (green). The RVxF motif of SHOC2 (GVAF) is shown as a sphere for the glycine and sticks for the side chain of valine, alanine and phenylalanine. **b** The interaction of the RVxF motif of SHOC2 (pink cartoon) with PP1CA (green cartoon) is shown. Side chains are shown as sticks with hydrogen bonds shown as black dashes. **c** Schematic representation of the SHOC2 RVxF–PP1CA interaction interface, as analyzed by PDBSum (http://www.ebi.ac.uk/pdbsum/). The interactions are colored using the following notations: hydrogen bonds as solid blue lines and non-bonded contacts as striped, orange lines (width of the lines is proportional to the number of atomic contacts). **d** Apparent K_D_ measurements of the SMP complex assembly for NS mutants and point mutants present at the RVxF motif. Noonan syndrome mutations are highlighted in black boxes. **e** Overall view of the SHOC2 LRR interactions (pink cartoon) with PP1CA (green surface). **f, g** Enlarged view of the N-terminal **(f)** and C-terminal **(g)** LRRs of SHOC2 with PP1CA as depicted in **e.** Side chains are shown in sticks and hydrogen bonds as black dashes. **h** Schematic representation of the SHOC2 LRRs–PP1CA interaction interface, as analyzed by PDBSum. Interactions are colored as described in **c** with the addition of salt bridges as solid orange lines. **i** NS mutations modeled onto the SMP structure. The double Q269H and H270Y mutation in SHOC2 increases contacts between Y270 of SHOC2 and I45 of PP1C relative to the wild-type H270. The P50R mutation of PP1CA (P49R as originally identified in PP1CB) would result in a *de novo* interaction with N225 of SHOC2 (shown as an electrostatic surface). A57P of PP1CA (A56P as originally identified in PP1CB) surrounds the residues that form the hydrophobic pocket that the RVxF motif interacts with. The E184A mutation of PP1CA (E183A as originally identified in PP1CB) relives the charge-charge repulsion with E155 of SHOC2.

Several NS mutations are present near the SHOC2-PP1CA interface, and structural analysis explains why they are gain-of-function mutations. Normally SHOC2 H270 interacts with I45 of PP1CA through van der Waals interactions. The SHOC2 NS double mutation Q269H/H270Y^17^ potentially forms larger contacts between I45 of PP1CA and Y270 of SHOC2 (Fig. 3i), which may increase the affinity and therefore the lifetime of the complex. Three NS mutations have been identified in PP1CB^43,44^ and these residues are conserved across all three isoforms. One of these, P50R (P49R in PP1CB), appears to form a *de novo* contact with SHOC2, potentially forming a hydrogen bond to the main chain of N225 in SHOC2 (Fig. 3i). This mutation increases the apparent affinity of SMP complex formation ∼4-fold to 33 nM (Fig. 3d and Supplementary Fig. 5b) with a much slower off-rate as determined by SPR. The NS mutation A57P (A56P in PP1CB) does not contact SHOC2 directly. A57 is found within a loop adjacent to the RVxF motif binding site of PP1CA, which may affect SHOC2 binding, however, we observed no difference in complex formation by SPR compared to wild type (Fig. 3d and Supplementary Fig. 5b). The NS mutation E184A (E183A in PP1CB) forms van der Waals interactions with two residues of SHOC2 (E155 and H178); however, the carboxylic acid groups of E184 and E155 of PP1CA are in close proximity to each other (Fig. 3i). This NS mutation relieves charge-charge repulsion resulting in a ∼4-fold increase in apparent affinity to 35 nM as measured by SPR (Fig. 3d and Supplementary Fig. 5b).

### The SHOC2-MRAS interface

The SHOC2-MRAS interface buries the second largest surface area (2000 Å^2^) in the SMP complex. MRAS binds to the concave surface of SHOC2, which is adjacent to the PP1CA-SHOC2 (LRR) interface (Fig. 4a). Specifically, MRAS contacts LRRs 1-10, 12 and 14-16 (Supplementary Fig. 6a-b). Residues in the switch II region of MRAS predominately contact the LRRs 1-4, with further contacts in LRRs 6-7. Switch II engages predominately through van der Waals interactions (Fig. 4b) with only one hydrogen bond forming between Q80 of MRAS and D106 of SHOC2. MRAS switch I residues, however, interact with SHOC2 LRRs 4-6 and 8-10, forming five hydrogen bonds and three salt bridges (Fig. 4b). In addition, several residues within the C-terminal region of MRAS interact with the C-terminal residues of SHOC2, specifically H132 of MRAS forms a hydrogen bond to E428 while K158 of MRAS forms a salt bridge with E406 of SHOC2 (Fig. 4c). In total, seven hydrogen bonds and six salt bridges are formed at the SHOC2-MRAS interface (Fig. 4d). A schematic of a single LRR is shown in Fig. 4e. The HCS motif (LxxLxLxxN(x)_1-2_L) forms the concave surface of SHOC2. All interactions of switch I and II residues of MRAS with LRRs 1-10 are with the variable residues preceding the conserved asparagine of the HCS motif (Lx<u>x</u>L<u>x</u>L<u>x</u>xN(x)_1-2_L, Fig. 4e). The residues that MRAS contact in the C-terminal half of LRRs of SHOC2 are the earlier variable residues in the HCS motif (L<u>xx</u>LxLxN(x)_1-2_L, Fig. 4e) and as such are found higher up on the molecular surface of SHOC2 (Fig. 4f). Three NS mutations arise in MRAS that are constitutively active variants, specifically, G23V, T68I and Q71R (Supplementary Fig. 6c), but none of these MRAS residues directly contact SHOC2 or PP1CA,^27,45,46^.

**Figure 4.**
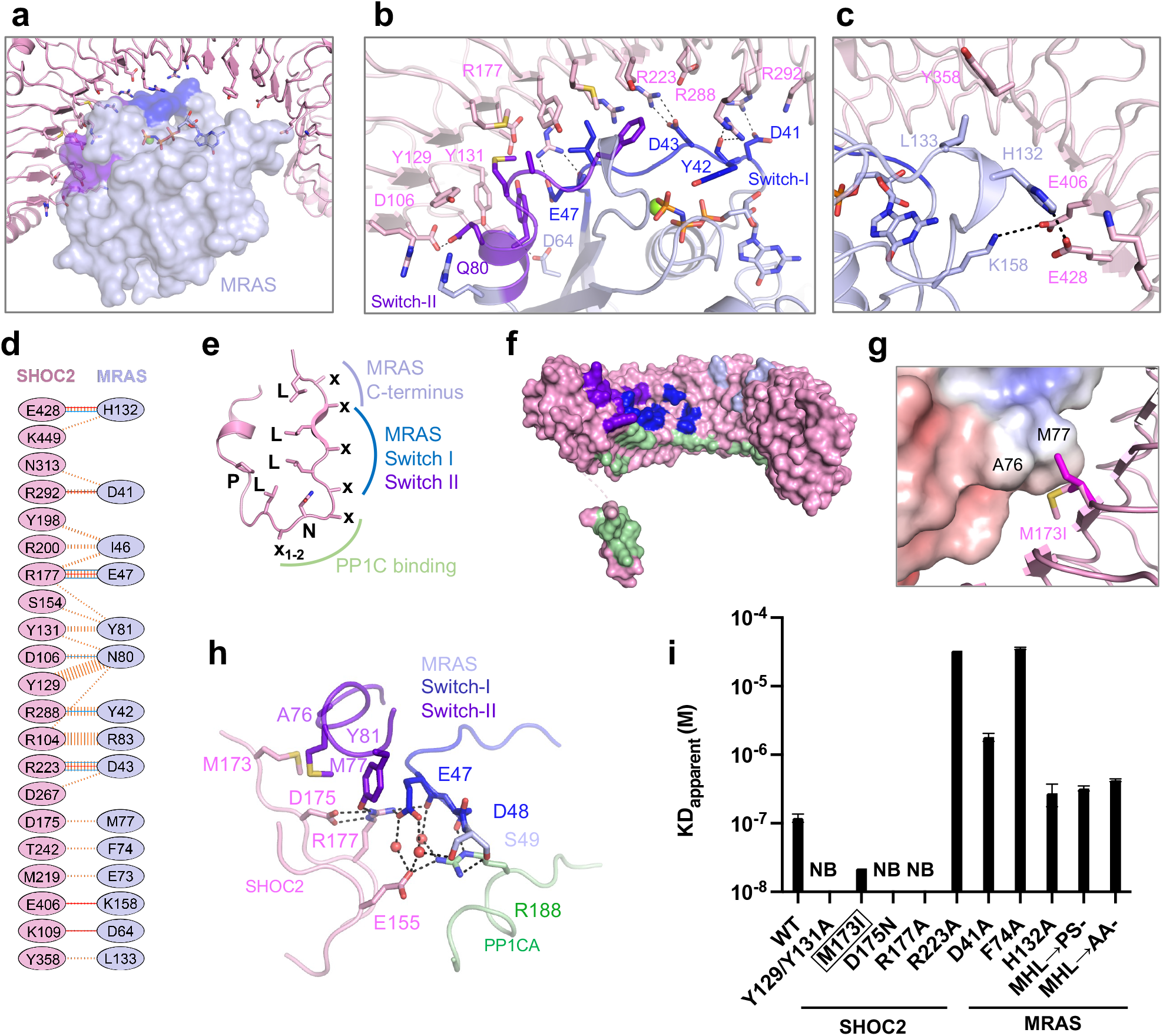
Structural and mutational analysis of the SHOC2-MRAS interface. **a** Overall view of SHOC2 (pink cartoon) interacting with MRAS (blue surface). Switch I, switch II, nucleotide and Mg^2+^ are shown in dark blue, purple, sticks and green sphere, respectively. **b, c** Enlarged view of the LRRs of SHOC2 interacting with switch I and switch II of MRAS **(b)**, and the C-terminus of MRAS **(c)**. **d** Schematic representation of the SHOC2-MRAS interaction interface, as analyzed by PDBSum (http://www.ebi.ac.uk/pdbsum/). The interactions are colored using the following notations: hydrogen bonds as solid blue lines, salt bridges as solid orange lines, and non-bonded contacts as striped, orange lines (width of the lines is proportional to the number of atomic contacts). **e** Schematic representation of a single LRR with the LRR sequence motif mapped onto it. Interactions of the LRRs with MRAS and PP1C occur through the top and midriff residues of SHOC2, while PP1C interacts through the bottom residues of SHOC2. **f** Surface of SHOC2 (pink) with residues contacted by switch I (dark blue), switch II (purple), and the C-terminus (blue) of MRAS and PP1CA highlighted. **g** The NS mutation M173I identified in SHOC2 is shown. M173 barely contacts A76 of MRAS, however, the NS M173I mutation potentially forms new, stronger interactions with M77 of MRAS. **h** A critical interaction of MRAS switch I (dark blue) and switch II (purple) with SHOC2 (pink) and PP1CA (green) either directly or through bridging waters (red spheres). **i** Apparent K_D_ measurements of the SMP complex assembly for NS mutants and point mutants present at the SHOC2-MRAS interface. No binding is indicated as NB. Noonan syndrome mutation is highlighted in a black box.

The SHOC2 mutation, M173I, was found in patients with overlapping Noonan and Cardio-Facio-Cutaneous syndromes^11,16^. SPR measurements of SHOC2 M173I show a 5-fold increase in the apparent affinity of SMP complexation (Supplementary Fig. 6d). Specifically, a slightly faster on-rate and a slower off-rate are observed relative to wild-type proteins demonstrating that this gain-of-function mutation stabilizes the complex. SHOC2 M173 does not contact MRAS. However, the substitution of methionine by isoleucine results in increased hydrophobicity and potentially forms a *de novo* contact with M77 of MRAS (Fig. 4g). SHOC2 D175 was identified as loss of function when mutated to asparagine in a genetic screen of soc-2, the SHOC2 homolog in *C. elegans*^12^. D175 forms a contact with M77 of switch II in MRAS. We believe the loss of function arises not from the contact with MRAS, but the removal of hydrogen bonds to the guanidino head group of R177 of SHOC2, which pre-orientates R177 to interact with MRAS (Fig. 4h). R177 contacts both switch I and II regions, specifically interacting with E47 and Y81, respectively. SHOC2_D175N_ or SHOC2_R177A_ mutations result in no complex formation (Fig. 4h-i and Supplementary Fig. 6d).

Additional mutations were made in SHOC2 and MRAS to identify key interactions at the SHOC2-MRAS interface. SHOC2 R223 interacts with D43 of switch I of MRAS, while SHOC2 Y129 and Y131 contact Q80 and Y81 in switch II of MRAS. SHOC2_R223A_ results in a weaker complexation as observed by SPR (∼300-fold reduction in the apparent K_D_), while no binding is observed for SHOC2_Y129A/Y131A_ (Fig. 4i and Supplementary Fig. 6d). MRAS D41 present in the switch I region interacts with R292 of SHOC2. The D41A mutation results in ∼10-fold weakening of the SMP complex compared to wild type (Fig. 4i and Supplementary Fig. 7a). MRAS F74 of switch II protrudes towards switch I and interacts with T242 of SHOC2. The loss of the phenyl group through the F74A mutation causes a ∼300-fold weakening of the complex (Fig. 4i and Supplementary Fig. 7a). The C-terminal residue of MRAS, H132, is found within a helical loop and forms a hydrogen bond to the E428 of SHOC2 only in the SMP1 complex where SHOC2 is pushed towards MRAS and PP1CA (Fig. 2d and Supplementary Fig. 2a). In K/H/NRAS, this helical loop is one residue shorter, and the histidine is replaced by shorter aliphatic residues, potentially resulting in the loss of this interaction (Supplementary Fig. 7b). Mutation of this MRAS residue results in a ∼3-fold weakening of complex formation (Fig. 4i and Supplementary Fig. 7a). Furthermore, mutation of this loop to its equivalent as observed in KRAS (^131^MHL^133^→PS-) or HRAS (^131^MHL^133^→AA-) also results in a similar 3-fold weakening of the SMP complex, suggesting that this region of MRAS contributes to higher affinity complexation over K/H/NRAS and that SHOC2 does interact with the region as observed in the SMP1 complex (Fig. 2d, 4i and Supplementary Fig. 7a).

### The MRAS-PP1CA interface

The MRAS-PP1CA interaction contributes the smallest buried surface to the complex with 1400 Å^2^ of buried surface area. MRAS binds to PP1CA adjacent to the PP1C-SHOC2 (LRR) interface in a way that all MRAS residues are > 20 Å away from the PP1CA active site (Fig. 5a). PP1CA contacts three different regions on MRAS (Fig. 5b); ***(i) N-terminal contacts*** – The first ten residues are unique to MRAS. These residues are not present in the classical RAS proteins (Supplementary Fig. 7b). Residues 4-6 of MRAS interact with PP1CA. This includes a hydrogen bond between S4 of MRAS and E218 of PP1CA and van der Waals interactions. These residues occupy the Myosin Phosphatase N-terminal Element (MyPhoNE) cleft on PP1CA. The myosin phosphatase targeting subunit 1 (MYPT1) protein uses an RVxF and MyPhoNE motif to bind to PP1CA. Although we observe MRAS occupying the MyPhoNE cleft, it does so differently compared to MYPT1 (Fig. 5c and Supplementary Fig 8a)^47^. Deletion of these N-terminal residues in MRAS results in a 6-fold weakening of SMP complexation as observed by ITC, confirming their importance for binding and as a key region of specificity between MRAS and the classical RAS proteins (Supplementary Fig. 8b). ***(ii) Pre-switch I contacts*** – residues 31-37 of MRAS interact within a pocket of PP1CA formed from the α_G_-α_H_ loop (residues 189-198) forming two hydrogen bonds and van der Waals interactions (Fig. 5b). ***(iii) Interswitch contacts*** – residues 48-53 of MRAS interact with PP1CA residues present in the β_6_-α_G_ loop and the α_G_-helix (residues 178-190). H53 of MRAS interacts with D179 of PP1CA. The MRAS H53A mutation weakens SMP complexation by ∼10-fold, as shown by SPR (Fig. 5d and Supplementary Fig. 8c). R188 of PP1CA is the only residue in the entirety of the SMP complex that engages with the other two proteins. Specifically, it forms a hydrogen bond to E155 of SHOC2, and D48 and S49 in MRAS (Fig. 4h, 5b). This potentially makes R188 the linchpin of the SMP complex. SPR measurements of SMP assembly with PP1CA_R188A_ show that no SMP complex is formed with this PP1CA mutant (Fig. 5d and Supplementary Fig. 8d). Despite being the smallest protein-protein interface, PP1CA and MRAS form eight hydrogen bonds and two salt bridges (Fig. 5e).

**Figure 5.**
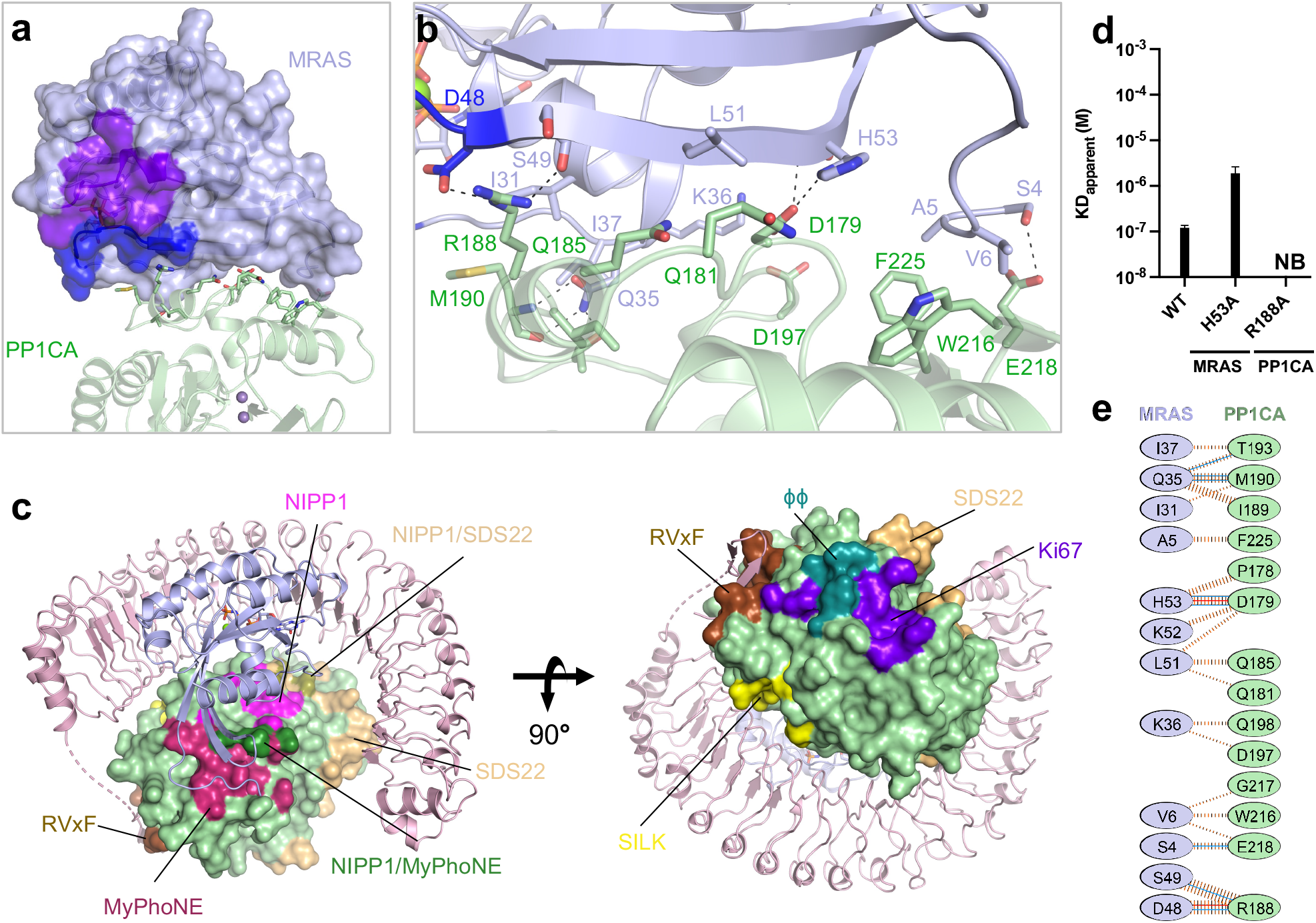
Structural and mutational analysis of the MRAS-PP1CA interface. **a** Overall view of PP1CA (green) interacting with the surface of MRAS (blue surface), with the switch I, switch II, nucleotide and Mg^2+^ shown in dark blue, purple, sticks and green sphere, respectively. The active site Mn^2+^ ions are shown as gray spheres. **b** Zoomed view of the PP1CA-MRAS interaction interface with side chains shown as sticks and hydrogen bonds as black dashed lines. **c** PP1CA is shown as a surface in the context of the SMP complex. All known PP1C interaction sites are colored on the surface of PP1CA (green) in the SMP complex; RVxF (brown), SILK (yellow), SDS22 binding site (wheat), LL (teal), ki67 binding site (purple), MyPhoNE (dark red), NIPP1 helix (magenta), overlap of MyPhoNE and NIPP1 helix (dark green), and overlap of the NIPP1 and SDS22 binding sites (olive). **d** Apparent K_D_ measurements of the SMP complex assembly for point mutants present at the MRAS-PP1CA interface. No binding is indicated as NB. **e** Schematic representation of the MRAS-PP1CA interaction interface, as analyzed by PDBSum (http://www.ebi.ac.uk/pdbsum/). The interactions are colored using the following notations: hydrogen bonds as solid blue lines, salt bridges as solid orange lines, and non-bonded contacts as striped, orange lines (width of the lines are proportional to the number of atomic contacts).

### Recognition of RAF substrates by the SMP complex

PP1CA has three active site channels/grooves denoted the acidic, the hydrophobic, and the C-terminal channels (Fig. 6a)^48^. To understand how the SMP recognizes RAF substrates, we tried to crystallize the SMP complex with RAF CR2-pS peptides, unfortunately, this was unsuccessful. We instead used the CABS-Dock server to dock a 15-mer BRAF CR2-pS peptide to predict which channels RAF substrates may use^49^. The majority of the top cluster containing 202 docked peptides were placed with the CR2-pS S365 in the active site with the N- and C-termini of the peptides occupying the acidic and hydrophobic channels, respectively (Fig. 6b). We observed similar docking poses with a 15-mer CRAF CR2-pS peptide (Supplementary Fig. 9a). Two NS mutations have been identified in the active site channels. D253Y (D252Y in PP1CB) and E275K (E274K) are found in the acidic and hydrophobic channels (Supplementary Fig. 9b). None of the top scoring peptides models contact these residues suggesting that these residues may selectively prevent other substrates from competing with RAF in the SMP complex or fine tune the affinity for RAF.

**Figure 6.**
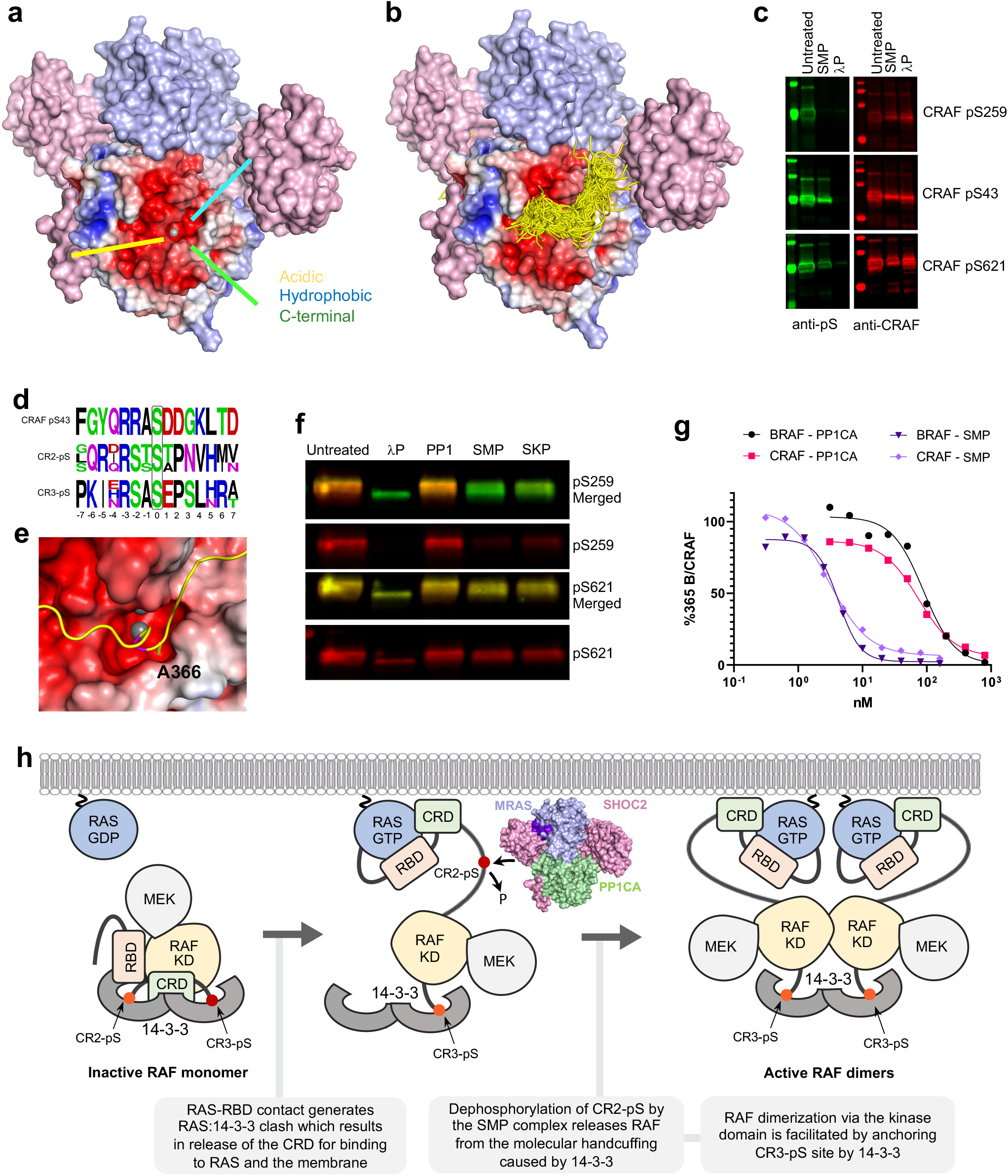
Model of recognition of RAF substrates by the SMP complex. **a** Structure of the SMP complex is shown as a surface. SHOC2 and MRAS are colored pink and blue, respectively. The surface of PP1CA is shown as an electrostatic surface as calculated by APBS. The three active site channels – acidic, hydrophobic and C-terminal are shown as yellow, cyan and green lines, respectively. Mn^2+^ ion is shown as a gray sphere. **b** The CABS-dock server was used to generate a 15-mer peptide of the CR2-pS region of BRAF and dock into the PP1CA structure of the SMP complex. All 202 peptides from the top cluster of solutions are presented as ribbons. The vast majority being placed in the active site, with all peptides placed with their N- and C-termini in the acidic and hydrophobic active site channels. **c** Fluorescent western blot of CRAF either untreated or treated with SMP or lambda phosphatase (λP). Right-hand panels show the total CRAF present (red), while left-hand panels reveal CRAF by specific phosphoserine antibodies (green) targeting pS259 (top), pS43 (middle), and pS621 (bottom). Lambda phosphatase removes all phosphates, while the SMP complex only removes pS259. **d** Sequence alignments of the CRAF pS43, CR2-pS of ARAF, BRAF and CRAF, and CR3-pS of ARAF, BRAF and CRAF. The phosphoserine in each case is boxed in black at position 0. **e** The top docked CR2-pS peptide of BRAF is displayed as a ribbon in the active site with the PP1CA surface shown in electrostatic surface representation. S365 of BRAF present in the active site is colored magenta. The docked model suggests that A366 of BRAF would be placed inside the narrow negatively charged active site channel. This residue is an aspartic acid in the pS43 of CRAF and a glutamic acid in the CR3-pS peptides, offering a possible reason for the selectivity of the SMP complex for CR2-pS phosphopeptides. **f** Fluorescent Western blot of CRAF either untreated or treated with λP, PP1CA, SMP or SKP. Phosphoserine-specific antibodies for pS259 and pS621 are shown in red. Total CRAF is shown in green. SMP and SKP complexes specifically dephosphorylate pS259 of CRAF.P, PP1CA, SMP or SKP. Phosphoserine-specific antibodies for pS259 and pS621 are shown in red. Total CRAF is shown in green. SMP and SKP complexes specifically dephosphorylate pS259 of CRAF. **g** Comparison of dephosphorylation activity (EC50) of PP1CA and SMP complex on BRAF and RAF substrates derived from Li-COR quantification of bands from Supplementary Fig. 6c. **h** Model showing the role of the SMP complex in the RAF activation process.

To validate the specificity of the SMP complex for various phosphorylation sites on B/CRAF, we performed dephosphorylation assays. Treatment of CRAF with lambda phosphatase non-specifically removes all phosphates (pS43, pS249, and pS621). However, treatment of CRAF with the SMP complex shows that it selectively dephosphorylates CR2-pS259 (Fig. 6c). Similar specific dephosphorylation of BRAF CR2-pS365 was observed when used as a substrate (Supplementary Fig. 9c-d). Examination of the sequence composition around different pS sites provides a rationale for CR2-pS site specificity. The CR2-pS site contains either threonine or alanine residues at the +1-position in RAF substrates (Fig. 6d). In contrast, the CR3-pS site in RAF substrates and the pS43 site in CRAF contain glutamic acid and aspartic acid, respectively, at this position (Fig. 6d). Docking of B/CRAF CR2-pS peptides shows that residues in the +1 position would be placed inside the restrictive negatively charged active site channel, suggesting a preference for small and non-acidic residues at this position (Fig. 6e). The +1-position of PP1C has already been noted to prefer to select against aspartic and glutamic acid residues in this position^50^. Dephosphorylation of a phosphorylated BRAF 15-mer CR2-pS peptide and one with the +1-position mutated to glutamic acid by the SMP complex was measured by MALDI-TOF. Results showed that the SMP complex readily dephosphorylates the wild-type BRAF CR2-pS peptide but is slower where the +1-position of the peptide is mutated to glutamic acid, which is consistent with our docking results (Supplementary Fig. 10).

Comparison of dephosphorylation using SKP and SMP complexes showed that the SKP complex has slightly weaker dephosphorylation activity for CRAF CR2-pS compared to the SMP complex (Fig. 6f). To determine if SHOC2 and MRAS have any effect on the dephosphorylation activity of PP1CA, we carried out dephosphorylation activity using apo-PP1CA and the SMP complex with BRAF and CRAF as substrates. Interestingly, the dephosphorylation activity of apo-PP1CA was 10-30-fold lower than the SMP complex yet still displays specificity to CR2-pS (Fig. 6g and Supplementary Fig. 9c-d) suggesting that MRAS and SHOC2 play a role in enhancing the dephosphorylation activity of PP1CA towards CR2-pS in RAF substrates.

## DISCUSSION

Here we present a high-resolution structure of the heterotrimeric SMP complex, which provides insights into how SHOC2, MRAS, and PP1CA interact to form this ternary complex and insight into RAF dephosphorylation. Analyses of NS mutants found in SHOC2, MRAS and PP1C in the SMP complex structure suggest how these substitutions would result in additional interactions resulting in tighter complex formation, sustained dephosphorylation of RAF, and activation of MAPK/ERK signaling. All three proteins form multiple contacts with each other but based on buried surface area upon complex formation and number of interactions, the SHOC2-PP1CA interface is most extensive. Interestingly, none of the three proteins form stable, high affinity, binary complexes with each other, highlighting the strikingly synergistic nature of SMP complex formation. We did observe a weak binary SHOC2-PP1CA interaction by SPR, suggesting this forms first. All PP1C regulators that rely on the RVxF motif to bind PP1C form high-affinity binary complexes. As SHOC2 contains an RVxF motif, it is unusual and unique to only observe a weak interaction with PP1C. Therefore, SHOC2 appears to be distinct from SDS22 (an LRR protein like SHOC2 but lacks an RVxF motif) and RVxF-containing proteins, both of which form high-affinity binary complexes with PP1C. Ternary complex assembly is only achieved with active MRAS, indicating that MRAS plays an important role in initiating and regulating the SMP complex assembly. As MRAS is anchored in the plasma membrane through its HVR, it targets PP1C to the plasma membrane via SHOC2. As MRAS only form complexes with PP1C in the presence of SHOC2, these data suggest that SHOC2 functions as an adaptor protein in this complex.

Both our and previous binding studies results suggest that K/H/NRAS can substitute for MRAS in the SMP complex^26^. However, *in vivo*, MRAS is most likely to form part of the SHOC2-RAS-PP1C complex for several reasons. Our SPR data shows a 7-40-fold higher affinity of complex formation with MRAS compared to K/H/NRAS, and that the SKP complex displays relatively weaker dephosphorylation activity compared to SMP. This increase in affinity observed with MRAS comes from the additional interactions from the N- and C-termini (residues 4-6 and H132) and compositional differences in interacting residues present in the pre-switch and interswitch regions of MRAS. Previous studies have shown that substitution of MRAS residues by corresponding residues in KRAS in the pre-switch-I and interswitch regions decreases MRAS affinity for SHOC2/PP1C^11^. Substitution of L51 of MRAS to Arg (R41 in KRAS) increased MRAS affinity to B/CRAF, whereas it decreased its affinity for SHOC2/PP1C, suggesting that MRAS and canonical RAS proteins evolved to play different roles during the RAF activation process. This is supported by the observation that MRAS is unable to activate RAF kinases to the same extent as canonical RAS proteins, and it is likely due to differences in the interswitch region that affects MRAS interaction with CRD of RAF proteins^7,11^. However, it must be noted that SHOC2, but not MRAS or PP1C, has been repeatedly identified in synthetic lethality CRISPR/Cas9 screens^19–22^. Furthermore, MRAS KO does not phenocopy SHOC2 KO in mice^51,52^. It is therefore possible that in the absence of MRAS, the lower affinity interaction of canonical RAS proteins for SHOC2 and PP1C complexation may be sufficient for CR2-pS RAF dephosphorylation *in vivo*. Similarly, it was recently shown using H/N/KRAS-less mouse embryonic fibroblasts that MRAS could substitute for classical RAS proteins for ERK activation by RAF inhibitors^27,53^. Thus, MRAS and canonical RAS lower affinity interactions for RAF and SHOC2-PP1, respectively, may be sufficient to provide redundancy in some contexts.

The SMP complex is responsible for dephosphorylation of CR2-pS sites and activation of RAF. Our results show that the interaction with MRAS and SHOC2 selectively enhance the dephosphorylation activity of PP1CA ∼20-fold against CR2-pS but not any other RAF phospho-sites, suggesting that MRAS and SHOC2 do play a role in targeting and enhancing dephosphorylation of CR2-pS by PP1CA. SHOC2 and/or MRAS may aid in the recruitment of RAF through several different mechanisms which have been observed in other PP1C and PP1C-interacting protein (PIP) complexes. Several PIPs contain extra domains which interact with substrates either directly or indirectly. PP1C interaction with muscle glycogen–targeting (G_M_) regulatory subunit via RVxF and φφ motif is an example that involves both direct and indirect substrate recruitment^54^. A carbohydrate-binding domain in G_M_ binds to glycogen or muscle-specific glycogen synthase (GYS1). When bound to glycogen, the PP1C-G_M_ complex is localized in the vicinity of phosphorylase *a*, a substrate for PP1C, which also interacts with glycogen. Glycogen, therefore, mediates the interaction between phosphorylase *a* and the PP1C-G_M_ complex. GYS1 is also a substrate for PP1C, and it binds directly to the carbohydrate domain of G_M_, recruiting the substrate to the holoenzyme. As both MRAS and SHOC2 binding to PP1CA enhances dephosphorylation of RAF substrates *in vitro*, it suggests that a direct substrate recruitment mechanism is used. As MRAS can be substituted for the classical RAS proteins in this complex, it is tempting to suggest that RAF substrates would be recruited by RAS proteins through the RAS-binding domain (RBD) and cysteine-rich domain (CRD) of RAF. However, RAS uses their pre-switch, switch I and interswitch residues to bind these two domains of RAF, which are buried by SHOC2 and PP1CA^7^. This would suggest that either another region of MRAS aids in the recruitment of RAF or SHOC2. In addition to the direct substrate mechanism, an indirect substrate recruitment mechanism could also occur at the plasma membrane. Both MRAS and KRAS share a similar C-terminal hypervariable region (HVR), lipidation profile, and are both found to co-localize within the disordered lipid regions of the plasma membrane^55^. Active KRAS would therefore bind and recruit RAF substrates both temporally and spatially with the active SMP complex at the disordered lipid regions of the plasma membrane.

PP1C forms complexes with over 200 PIPs that bind to PP1C through short linear motifs (SLIMs) that dock to surface grooves of PP1C. The best characterized SLIM is the RVxF motif, which is present in the majority of PIPs but does not influence enzymatic activity as it is located away from the active site. Other SLIMs include SILK, MyPhoNE, φφ and SpiDoC motifs (Fig. 5c). PIPs use combinations of these SLIMs to form multivalent interactions with PP1C that enhance regulator binding avidity and create PP1C holoenzymes with unique properties and substrate specificity, although the exact molecular mechanisms how they alter substrate specificity are unclear. This is true for the SMP complex as both SHOC2 and MRAS bind >20 Å from the active site. We do not observe any alteration of residues or electrostatics of the active site channels of PP1CA upon complexation, or extension of these active site channels as seen in the Phatcr1-PP1CA complex^56^. However, the entrance to the acidic channel may be partially blocked due to the disordered residues between the RVxF motif and LRRs of SHOC2 as seen in the NIPP1-PP1CA complex^57^. The formation of the SMP complex at the plasma membrane is likely to prevent the formation of other PP1C-holoenzymes due to SHOC2 and MRAS occluding several PIP-binding sites on PP1C, including the RVxF, SILK, SDS22, MyPhoNE, and NIPP1 helix binding pockets (Fig. 5c)^29,31,58^.

The RAF activation cycle starts when active RAS interacts with RBD in the autoinhibited RAF complex (Fig. 6h). The RAS–RAF RBD interaction causes a steric clash between RAS and 14-3-3, resulting in conformational changes that dislodge the RBD and CRD from the autoinhibited RAF complex. This action allows the CRD to interact with the plasma membrane and RAS to further stabilize the RAS–RAF interaction (Fig. 6h). The release of the CRD exposes the CR2-pS site. Dephosphorylation of this pS site by the SMP complex allows the exposed kinase domain to dimerize, forming an active dimeric RAF complex, stabilized by binding a 14-3-3 dimer to the CR3-pS sites (Fig. 6h). Our *in vitro* assays and previous studies show that the SMP can dephosphorylate RAF without RAS though this would not happen inside the cell. The membrane-bound SMP complex would not dephosphorylate RAF unless it is recruited to the plasma membrane by active RAS.

The SMP complex is a high-value target for regulating RAF and MAPK-pathway activation. Considering SHOC2, MRAS and PP1C do not form a high-affinity binary complex and several interface mutants described in this study disrupt complex formation, targeting any of the three interaction interfaces would likely disrupt the SMP complex formation. The MRAS switch regions interact extensively with SHOC2, and any compound that binds to the switch-II pocket of MRAS could prevent SMP complex formation, although the compound would have to be able to bind other RAS proteins to prevent their substitution for MRAS. However, any such compound would inhibit multiple RAS effectors and thus will lack specificity. Targeting PP1C, specifically in the context of the SMP complex, would potentially be difficult due to the large number of proteins that bind to PP1C (Fig. 5c). Targeting the RVxF pocket on PP1C for example would prevent the majority of PIPs from binding, however it may be possible to target PP1C selectively. Our results show that R188 of PP1C acts as a linchpin in the SMP complex as it is the only residue that contacts the other two proteins. R188 is not part of any SLIM, so disrupting R188 with a small molecule could prevent SMP complexation specifically. Although the biology of SHOC2 is the least understood of the three proteins, SHOC2 does make an interesting target due to its identification in several synthetic lethality CRISPR/Cas9 screens, though it is unknown whether the loss of the SMP complex causes the lethality or the loss of a different SHOC2 interaction. Based on other SHOC2 studies, it appears that the MRAS binding region on SHOC2 is a unique interaction site and thus a promising druggable site^59,60^. Our data support that altering the surface of SHOC2 in LRR2 and LRR4 (Y129/131A or R177A, respectively) prevent the formation of the SMP complex through disruption of the SHOC2-MRAS interface. LRR4 also contains the loss-of-function mutation D175N and the NS mutation M173L, suggesting these two regions of the SHOC2 surface could be exploited as druggable target sites. The question as to how the SMP complex interacts and dephosphorylates the RAF/RAS complex remains unanswered. A deeper understanding of how these two complexes interact with each other at the plasma membrane could lead to new approaches to target RAS/RAF-driven cancers and Noonan syndrome.

## METHODS

### Purification of recombinant proteins

Proteins for crystallography, human SHOC2_58-564_ and the SMP complex, were cloned, expressed, and purified as previously described^36^. Proteins for SPR and phosphatase assays were generated from DNA constructs initially synthesized as Gateway Entry clones (ATUM, Newark, CA). Constructs consisted of gene optimized fragments containing an upstream tobacco etch virus (TEV) protease site (ENLYFQ/G) followed by the coding sequence of human PP1CA (amino acids 7-300), human MRAS (amino acids 1-178), or human SHOC2 (amino acids 2-582) with mutations described in the results. Constructs were optimized for expression in *E. coli* (PP1CA, PP1CB and MRAS) or insect cells (SHOC2). Entry clones were transferred to *E. coli* or baculovirus expression clones containing amino-terminal His6-MBP (maltose-binding protein) fusions by Gateway LR recombination (Thermo Fisher Scientific, Waltham, MA) into pDest-566 (*E. coli*, Addgene #11517) or pDest-636 (baculovirus, Addgene #159574). Final baculovirus expression clones were used to generate bacmid DNA in strain DE95 using the Bac-to-Bac system (Thermo Fisher Scientific, Waltham, MA).

### Protein expression and purification

MRAS proteins were expressed as described for the Dynamite expression protocol^61^. PP1CA proteins were expressed in a similar manner but with some changes. Specifically, the expression strain also included the GroEL-expressing plasmid pG-tf2 (Takara Bio USA, Inc.), and expression was induced at 10°C. MRAS proteins were purified essentially as outlined in Kopra *et al*. for KRAS (1-169) with 1 mM MgCl_2_ in the final buffer^62^. PP1CA proteins were purified with modifications to the approach outlined for KRAS (1-169)^62^. Specifically, the lysis buffer was 20 mM Tris-HCl, pH 8.0, 700 mM NaCl, 10% glycerol (w/v), 1 mM MnSO_4_ (or MnCl_2_), 1 mM TCEP, and 0.5% Triton X-100 (w/v), the same buffer without Triton X-100 was used in subsequent steps until the SEC/final buffer, which was 20 mM Tris-HCl, pH 8.0, 500 mM NaCl, 1.0 mM MnSO_4_, and 1 mM TCEP, clarification of the lysate required extended conditions to overcome the presence of glycerol in the buffer (2 hours at 13,000 x *g*), and a 5 ml MBPTrap HP column (Cytiva, Marlborough, MA) was placed in front of the preparative SEC column to capture undigested fusion protein. All mutant protein SEC elution profiles and measured thermal denaturation temperatures were similar to the wild-type proteins.

### Nucleotide Exchange

MRAS-GDP (the protein is normally in the GDP-bound state when purified from *E. coli*) is mixed with a 5-molar excess of non-hydrolysable GMPPNP (tetralithium salt, Jena Biosciences NU-401-50) in a reaction mixture of 200 mM ammonium sulfate and 100 µM ZnCl_2_. The final MgCl_2_ concentration in the reaction is less than 1 mM through dilution of the stock protein with the reaction mixture components. The typical protein concentration range in the reaction was 0.1–0.3 mM. Alkaline phosphatase-agarose beads (Sigma P0762-250UN) were added at a ratio of 1U per mg of protein and the reaction was mixed at room temperature for 3 hr. The beads were then removed by centrifugation at 1500 x *g* for 2 min. The sample was adjusted with an additional 10-fold molar concentration of GMPPNP and incubated at 4°C for two hours or overnight. Excess nucleotide was removed by passing over a PD-10 desalting column packed with Sephadex G-25 resin (Cat # 17085101, Cytiva, Marlborough, MA) in 20 mM HEPES, pH 7.4, 150 mM NaCl, 1 mM MgCl_2_, and 1 mM TCEP. Protein concentration was determined on a Nanodrop 2000C spectrophotometer (Thermo Fisher Scientific, Waltham, MA) reading at A_280_.

### Crystallization and data collection

Purified SMP complex was concentrated to 15 mg/ml, and crystallization screening was carried out at 20□°C using the sitting-drop vapor diffusion method by mixing purified SMP complex with an equal volume of reservoir solution (200 nL:200 nL). Crystals of the SMP complex appeared within 24 hours in the crystallization condition containing 25% w/v PEG 1500, 0.1 M MMT pH 4.0. These crystals, cryoprotected with 20% v/v of glycerol, diffracted anisotropically to a resolution of ∼3.7 Å. To improve the diffraction quality of these crystals and the stability of the SMP complex during the crystallization, GTP present in the MRAS of the SMP complex was exchanged with GMPPNP. Further optimization of the crystallization condition was carried out by increasing the pH (0.1 M MMT, pH 4.2) and reducing the concentration of PEG 1500 (15% w/v). However, these optimized crystals only diffracted to 3.2 Å and remained anisotropic. Matrix micro-seeding was performed to further improve the quality of diffraction^63^. Briefly, two drops worth of SMP crystals were transferred to a seed bead tube (Hampton Research) containing 100 μL of 15% PEG 1500, 0.1M MMT, pH 4.2, vortexed for 30 s, before dilution to 1 mL with 15% PEG 1500, 0.1M MMT, pH 4.2. Another round of extensive crystallization screening was carried out in which a ratio of 200 nL protein:133 nL reservoir:67nl of seeds was used. Approximately 30 new conditions were identified, though only one yielded isotropic diffracting crystals to 2.8 Å (20% w/v PEG 3350, 0.2M sodium sulfate). Crystals were further optimized around this crystallization condition through a grid screen and seeding. The grid varied the concentration of PEG 3350 and sodium sulfate from 15-25% w/v (in steps of 1.43%) and 0-250 mM (in steps of 23 mM), respectively. A ratio of 200 nL protein:133 nL reservoir:67nl of seeds was used. Seeds were prepared fresh, as described above, using crystals from the original condition of 15% PEG 1500, 0.1M MMT, pH 4.2 (frozen seeds failed to work). A 2.17 Å dataset was collected on beamline 24-ID-C at the Advanced Photon Source (Argonne) with a crystal grown from 17.9% w/v PEG 3350, 136 mM sodium sulfate, and 1:10 dilution of seeds in a ratio of 200 nL protein:133 nL reservoir:67nl of seeds. The crystal was cryoprotected with 25% (v/v) glycerol.

To solve the structure of SHOC2, we carried out crystallization screening of two SHOC2 constructs (2-584; 58-564) using commercial screens at 15 mg/ml protein concentration. The SHOC2 construct ranging from 58-564 produced crystals in multiple ammonium sulfate conditions at low pH. Optimization of SHOC2 (58-564) crystals produced diffracting crystals in 1.5M ammonium sulfate, 0.1M sodium citrate pH 5.0. Crystals were cryoprotected with 30% glycerol, and a 2.4 Å dataset was collected on beamline 24-ID-C at the Advanced Photon Source (Argonne).

### Structure determination and analysis

Crystallographic datasets were indexed and integrated using XDS^64^. The integrated data were then scaled, truncated, and converted to structure factors using the program Aimless present in the CCP4 suite^65,66^. Matthew’s coefficient suggested two copies of the SMP complex inside the asymmetric unit. The structure was determined using the molecular replacement program Phaser using mouse MRAS bound with GMPPNP (PDB ID 1X1S) and human PP1CA (PDB ID 6DNO)^67^. This helped in locating two copies of MRAS and PP1C inside the asymmetric unit. Since the structure of SHOC2 was not available at this time, we used a Rosetta-generated model of SHOC2 as a search model in our molecular replacement runs^67^. Although this approach did not work, the initial maps calculated after placing two copies of MRAS and PP1C allowed the manual placement of the Rosetta-generated model of SHOC2. This was followed by a rigid body refinement. The initial model of the SMP complex was iteratively rebuilt in COOT and refined with Refmac5, followed by Phenix.Refine^66–69^. During the final stages of model building and refinement, water molecules were identified by the automatic water-picking algorithm in COOT and Refmac5/Phenix.refine. The positions of these automatically picked waters were checked manually during model building. The structure of SHOC2 was determined using SHOC2 present in the SMP complex as a search model in the molecular replacement Phaser^67^. This search identified one copy of SHOC2 in the asymmetric unit. Model building and refinement of SHOC2 were carried out using the same protocol as described above for the SMP complex. Secondary structural elements were assigned using DSSP (https://swift.cmbi.umcn.nl/gv/dssp/). Figures were generated with PyMOL, and surface electrostatics were calculated with APBS^70,71^. Crystallographic and structural analysis software support was provided by the SBGrid Consortium^72^. Data collection and refinement statistics are shown in Supplementary Table 1.

### SPR measurements

CM5 chips (Cytiva Life Sciences) were preconditioned by injecting 0.5% SDS, 100 mM HCl, 0.85% H_3_PO_4_, and 50 mM NaOH in that order at 30 µL min^-1^ for 60 seconds in PBS pH 7.4 running buffer. 200 µg/mL Neutravidin (Thermo Scientific) in 10 mM sodium acetate, pH 4.5 was amine coupled to the surface in PBS running buffer using standard EDC/NHS chemistry to a density of ∼7000 RU per flow cell. All buffers were vacuum filtered through 0.2 µm cellulose acetate membranes. Avi-tagged SHOC2 proteins were biotinylated *in vitro* using the procedure described previously^73^ and then captured by manual injection to an appropriate density in 10 mM HEPES, 150 mM NaCl, 2 mM MgCl_2_, 0.05% Tween 20, 1 mM TCEP, pH 7.4. Protein analytes MRAS and PP1CA were diluted equimolar to the highest concentration, typically 1 µM, in the buffer above then serially diluted three-fold four times in the same buffer for a total of five concentrations. Single-cycle kinetic responses consisted of injections at 30 µL min^-1^ with a contact time of 180 seconds and a dissociation time of 1600 seconds for each concentration of analytes. Sensorgrams were double referenced by subtracting the signal from a reference channel of neutravidin alone and a buffer blank. The data was fit to a 1:1 kinetic model to calculate an apparent K_D_ using the S200 evaluation software package, or the Insight software package. All experiments were conducted at 25 °C on an S200 or 8K instrument (Cytiva Life Sciences). All binding data are tabulated in Supplementary Table 2 with the number of replicates indicated. Errors were calculated from multiple experiments. Certain mutants could not be tested by SPR due to non-specific binding to the reference channel. In these cases, affinity measurements were performed by ITC.

### Isothermal titration calorimetry measurements

Proteins were extensively dialyzed against 30 mM HEPES, 500 mM NaCl, 1 mM MgCl_2_, 0.5 mM TCEP, 0.1 mM MnCl_2_, pH 7.5. Duplicate ITC measurements were performed on a MicroCal PEAQ-ITC instrument (Malvern Panalytical). An ITC experiment consisted of 15 µM of PP1C and MRAS in the cell with 175 µM of SHOC2 in the syringe. All measurements were carried out at 25°C, with a stirring speed of 750 rpm and 19 injections of 2 µl injected at 210s intervals. Data analysis was performed using a “one set of sites” model using the MicroCal PEAQ-ITC analysis software (Malvern Panalytical). All binding data are tabulated in Supplementary Table 3 with the number of replicates indicated.

### RAF kinase dephosphorylation assay by Western blotting

PP1CA and SMP complex phosphatase activity was tested on purified His6-CRAF protein, GST-CRAF or GST-BRAF. For His6-CRAF protein, substrate was diluted in 20 mM HEPES pH 7.4, 150 mM NaCl, 1 mM TCEP, 2 mM MgCl_2_, 2 mM MnCl_2_ to a final concentration of 686 nM. 20 µl of diluted CRAF/BRAF sample was mixed with 20 µl of 204 nM SMP or PP1CA and incubated at 30°C for 30 minutes. After 30 minutes, 40 µl of 2x NuPAGE LDS sample buffer (Thermo Scientific, Waltham, MA) was added to the tube, and samples were boiled for 5 minutes to stop the reaction. Western blots were prepared by electrophoresing 10 µl of each sample on an SDS-PAGE gel, transferring samples to a PVDF membrane via iBlot (Thermo Scientific, Waltham MA) using standard manufacturer’s conditions, and probing for CRAF pS43 (Abcam #ab150365), pS259 (Abcam #ab173539), pS621 (Abcam #ab4767), and anti-His6 for total CRAF (Abcam #ab18184). For GST-CRAF and GST-BRAF proteins, 50 nM of the substrate was incubated with 2-fold dilutions of either PP1CA (800-3.1 nM) or SMP complex (160-0.31 nM) for 1 hour at 37°C. Reactions were stopped by mixing with 4x LDS sample buffer. Western blots were prepared as described above and probed using antibodies against pS365 BRAF (in-house antibody), pS259 CRAF (CST #9421), BRAF F7 (SC #5284), CRAF (BD #610152), SHOC2 (in-house antibody) and PP1CA E-9 (SC #7482). Final images were taken using an Odyssey CLx (LI-COR Biosciences, Lincoln NE).

### RAF substrate docking

The CABS-Dock web server (http://biocomp.chem.uw.edu.pl/CABSdock) was used to dock BRAF and CRAF CR2-pS 15-mer peptides^49^. Briefly, ten peptides of RAF substrate are generated from a generic library and placed randomly approximately 20 Å from the surface of PP1CA. Each peptide undergoes 50 annealing cycles of a Replica Exchange Monte Carlo Scheme. Snapshots (1000) are taken of the trajectory of each starting peptide, resulting in 10000 initial models. Non-binding peptide models are removed and then sorted by calculating their protein-peptide interaction energy. The lowest 10% (1000 models, CA atoms only) are then clustered in a k-medoids procedure (k=10). RMSD of peptides in each cluster are then calculated. RMSD and cluster size are used as ranking parameters. The top model of each cluster is reconstructed to an all-atom complex using MODELLER.

### BRAF CR2-pS dephosphorylation assay by MALDI-TOF

SMP complex phosphatase activity was tested on two synthesized 15-mer phosphopeptides (Genscript) of BRAF (the N-terminus was acetylated). One peptide was of the wild type sequence (GQRDRSSpSAPNVHIN), while the second was mutated in the +1-position to glutamic acid (GQRDRSSpS<u>E</u>PNVHIN). Stock solutions of each peptide were made in water (∼10 mM). A 50 μl reaction of 600 μM peptide with 100 nM of the SMP complex diluted in 20 mM HEPES, 150 mM NaCl, 1 mM TCEP was carried out. Time points (2 μl) were taken at t= 0 h, 2 h and 16 h and mixed with 10 μl saturated sinapinic acid solution (10% acetonitrile, 0.1% TFA) and spotted onto 384 well sample MALDI-MS plate and allowed to air dry. Mass spectrometry covering the range 1500-2500 Da was carried out using a Bruker rapidfleX MALDI Tissuetyper in reflector mode with 2000 laser shots per spectrum.

### Data availability

The atomic coordinates and structure factors of the SMP complex and SHOC2 have been deposited in the Protein Data Bank and are available under accession numbers 7TVF and 7TVG, respectively.

## Acknowledgments

We thank Bill Gillette, Hannah Ambrose, Julia Cregger, Peter Frank, Brianna Higgins, Min Hong, Jennifer Mehalko, Ashley Mitchell, Shelley Perkins, Nitya Ramakrishnan, Mukul Sherekar, Matt Smith, Troy Taylor, Vanessa Wall, and Stephanie Widmeyer of the Protein Expression Laboratory (Frederick National Laboratory for Cancer Research) for their help in preparing recombinant proteins. We are grateful to Srisathiyanarayanan Dharmaiah and Daniel Czyzyk for their help with crystallization and Timothy Waybright for nucleotide analysis. We thank Michael Murphy (Cytiva Life Sciences) for his advice on fitting the SPR kinetic data. We thank Lucy Young for critical feedback on the manuscript. We acknowledge Wolfgang Peti for advice on the expression and purification of PP1CA. X-ray diffraction data were collected at the Northeastern Collaborative Access Team beamlines (24-ID-C/E), funded by the U.S. National Institutes of Health (NIGMS P30 GM124165). The Pilatus 6M detector on the 24-ID-C beamline is funded by an NIH-ORIP HEI grant (S10 RR029205). This research used resources of the Advanced Photon Source, a U.S. Department of Energy (D.O.E.) Office of Science User Facility operated for the D.O.E. Office of Science by Argonne National Laboratory under Contract DE-AC02-06CH11357. This project was funded in part with federal funds from the National Cancer Institute, National Institutes of Health Contract HHSN261200800001E. The content of this publication does not necessarily reflect the views or policies of the Department of Health and Human Services, and the mention of trade names, commercial products, or organizations does not imply endorsement by the U.S. Government.

## Author contributions

D.A.B. and D.K.S. carried out crystallography work, structural analysis, and ITC experiments; P.A. and A.G.S. performed S.P.R. measurements; N.H., M.D., D.E., and P.R-V. carried out enzymatic assays and Western analysis. K.S., M.D., S.M., and D.E. prepared recombinant proteins. L.I.F., D.V.N., P.R-V., and F.M. contributed to the structural and functional analysis. D.A.B. and D.K.S. wrote the manuscript with inputs from all co-authors.

## Competing interest statement

F.M. is a consultant for Amgen, Daiichi Ltd., Frontiers Med, Exuma Biotech, Ideaya Biosciences, Kura Oncology, Leidos Biomedical Research, Inc., PellePharm, Pfizer Inc., P.M.V. Pharma, and Quanta Therapeutics. F.M. is a consultant and co-founder for (with ownership interest including stock options) BridgeBio, Olema Pharmaceuticals, Inc., and Quartz. F.M. has received research grants from Daiichi Sankyo and Gilead Sciences and has a current grant from Boehringer-Ingelheim.

## Notes

https://www.rcsb.org/structure/7TVF

https://www.rcsb.org/structure/7TVG

