## Supplementary Material for "Structure of the SHOC2–MRAS–PP1C complex provides insights into RAF activation and Noonan syndrome"

### Supplementary Information

- Supplementary Figures 1-10
- Supplementary Tables 1-3

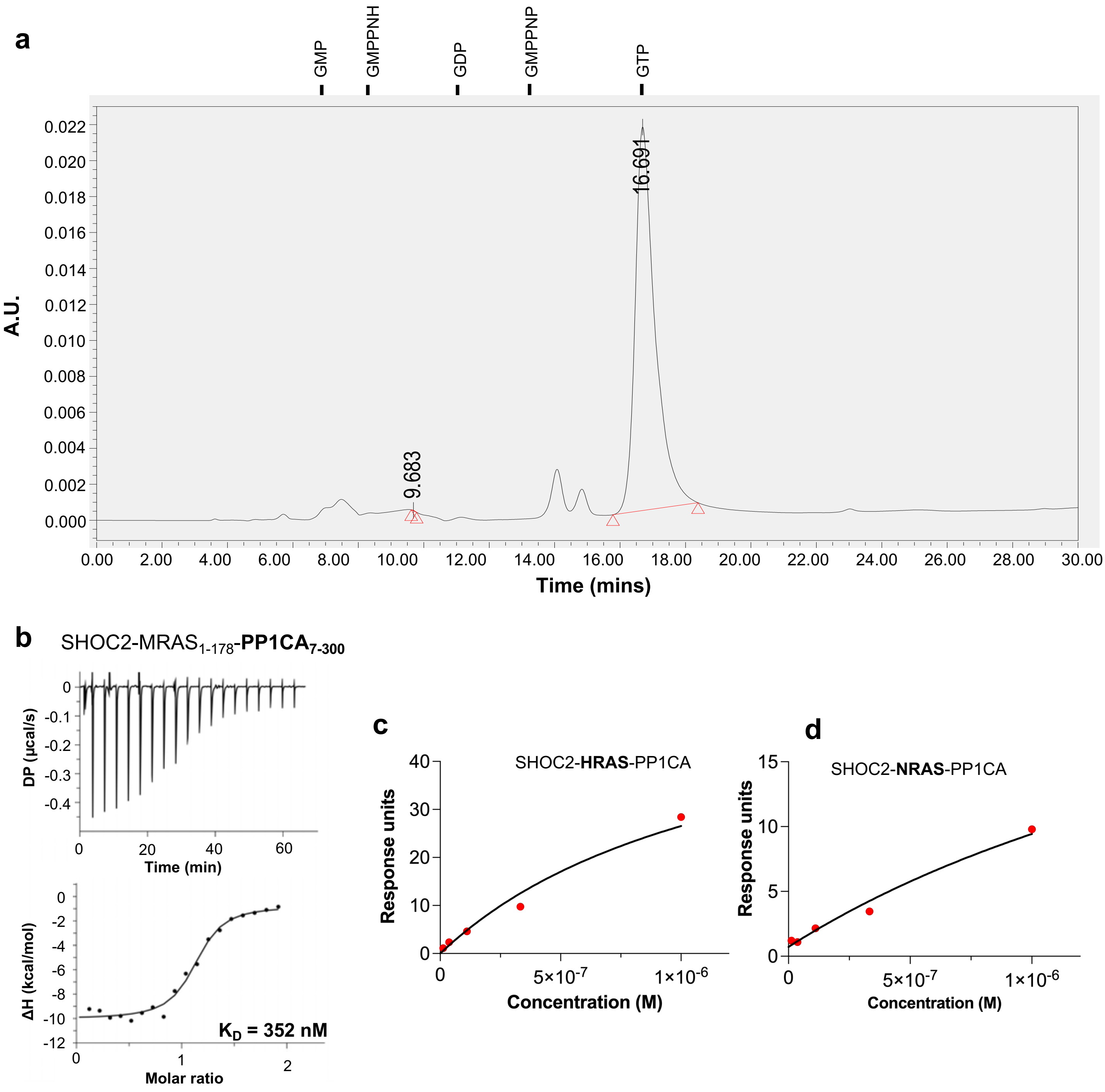

**Supplementary Figure 1. Assembly and selectivity of the SMP complex. a** Nucleotide analysis of the SMP complex by HPLC. Nucleotide standards and their retention time are shown above. **b** ITC experiment to measure the dissociation constant between SHOC2, $MRAS_{1\text{-}178}$ and $PP1CA_{7\text{-}300}$ **c** A steady-state plot of measured RU values from the formation of the SHOC2-HRAS-PP1CA complex against concentrations of $HRAS_{GMPPNP}$. **d** A steady-state plot of measured RU values from the formation of the SHOC2-NRAS-PP1CA complex against concentrations of $NRAS_{GMPPNP}$.

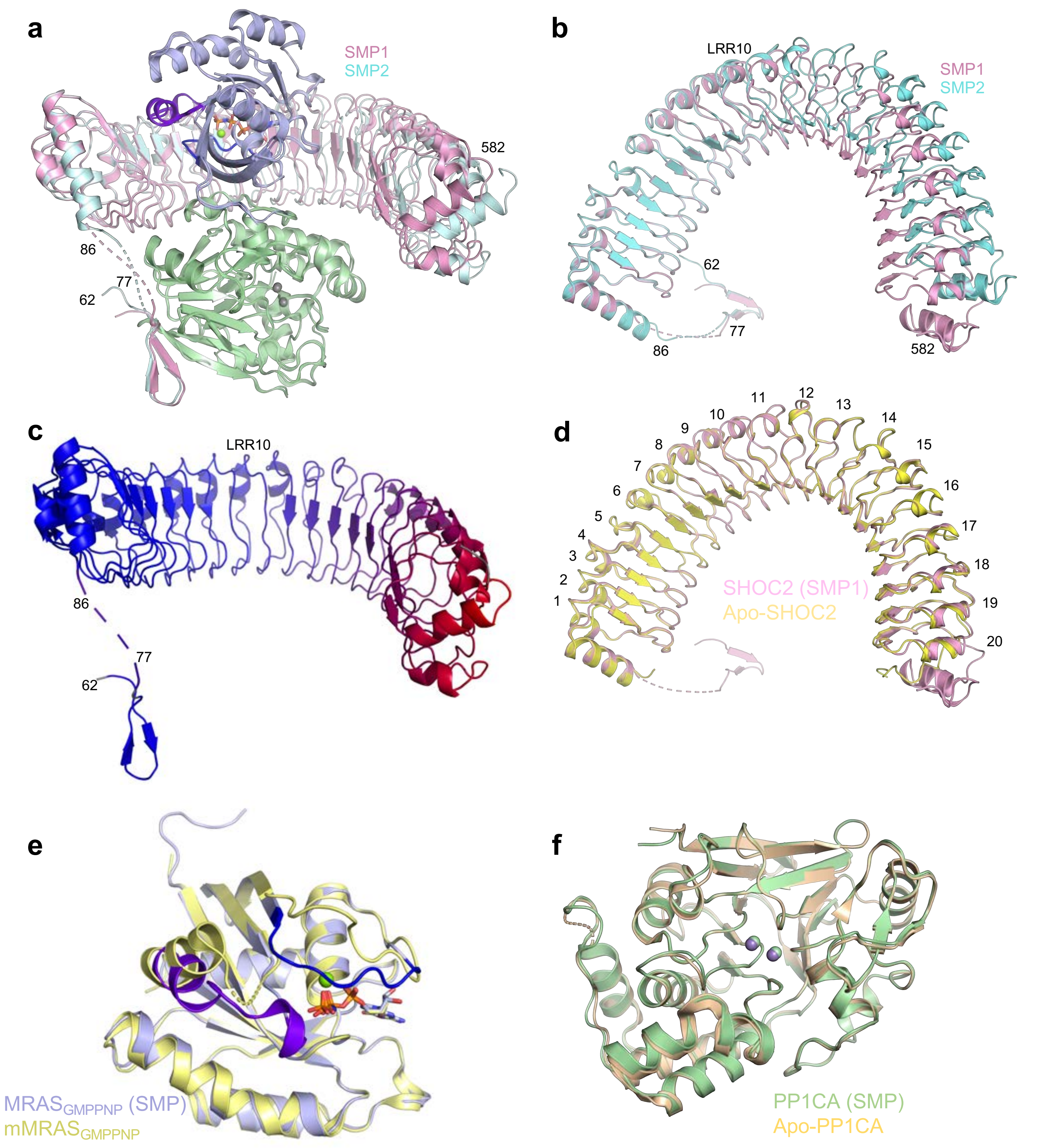

**Supplementary Figure 2. Comparison of the individual components of the SMP complex with their apo-forms. a** Superposition of the two SMP complexes found in the asymmetric subunit in cartoon form. Both chains of MRAS and PP1CA are in the same color, while the two SHOC2 chains are colored pink and cyan. The overall, SHOC2, MRAS and PP1CA RMSDs are 0.62 Å, 1.74 Å, 0.19 Å and 0.15 Å, respectively. **b** Top view as shown in panel **a** without MRAS and PP1CA present. LRR10 is marked, highlighting the hinge. **c** A cartoon of SHOC2 with a color gradient from blue to red showing the RMSD between the two SHOC2 molecules in the SMP complex, with blue and red representing low and high RMSD, respectively.

LRR10 is marked highlighting the hinge. **d** Superposition of apo-SHOC2 (yellow) with SHOC2 from the SMP complex (pink) which was used for all subsequent analysis. All LRRs are labeled. **e** Superposition of mouse MRAS bound to GMPPNP (yellow; PDB ID 1X1S, Ye, M. *et al*., (2005). J. Mol. Chem. 280: 31267-31275)) with human MRAS from the SMP complex (blue). Switch I, switch II, nucleotide and $Mg^{2+}$ ions are shown in dark blue, purple, sticks and green spheres, respectively. The overall RMSD is 0.32 Å. **f** Superposition of apo-PP1CA (olive, PDB ID 4MOV, Choy, M.S. *et al*., (2014). Proc. Natl. Acad. Sci. 111: 4097-4102) with PP1CA from the SMP complex (green). $Mn^{2+}$ ions from SMP and apo-PP1CA are shown in green and gray, respectively. The overall RMSD is 0.28 Å.

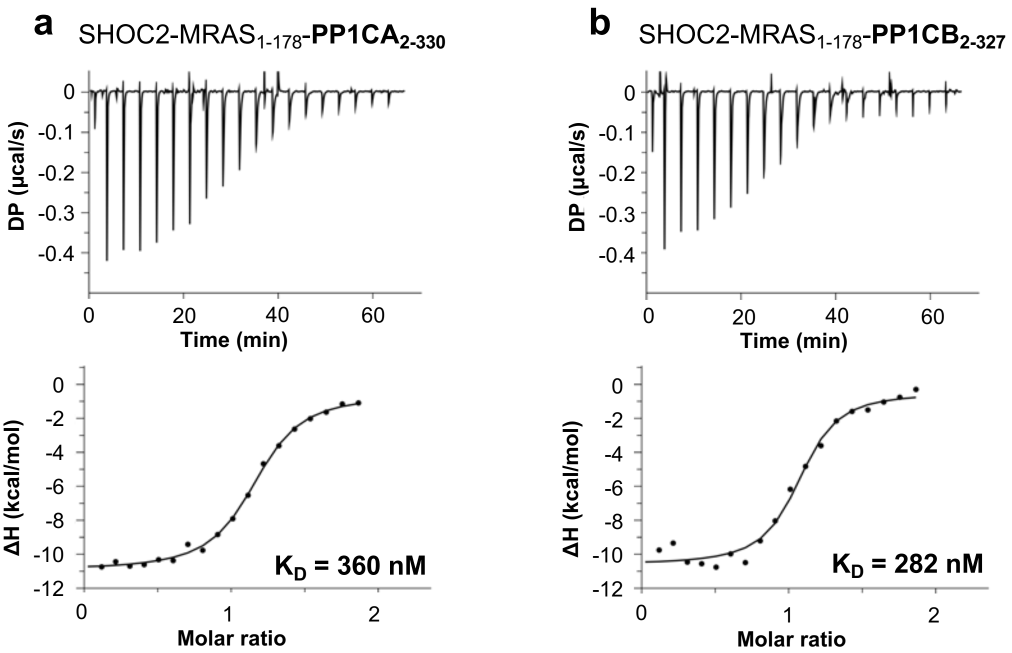

**Supplementary Figure 3. PP1C isoform specificity of the SMP complex. a, b**. ITC experiments to measure the dissociation constant between SHOC2, $MRAS_{1\text{-}178}$ and $PP1CA_{2\text{-}330}$ (a), and SHOC2, $MRAS_{1\text{-}178}$ and $PP1CB_{2\text{-}327}$ (b).

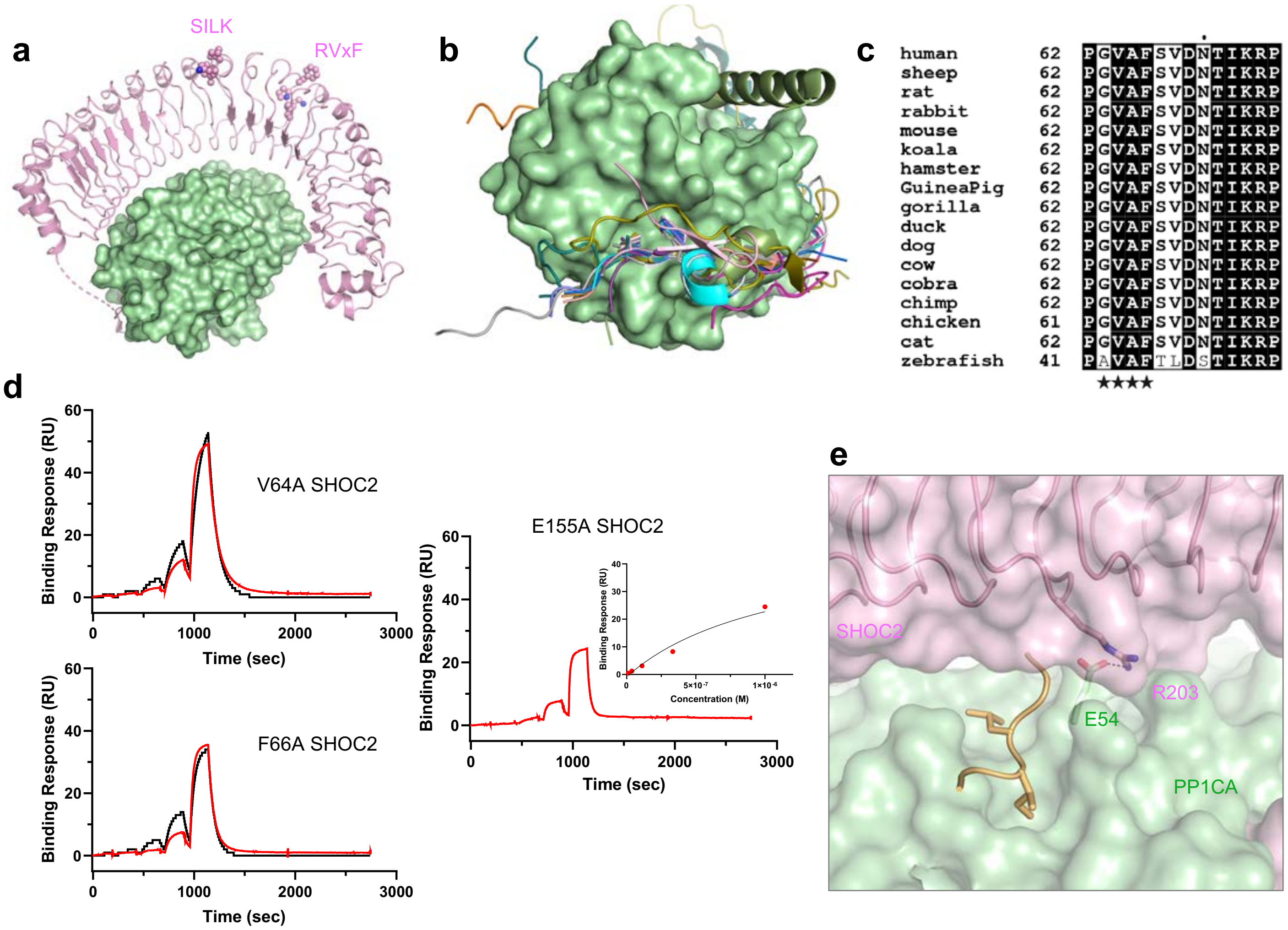

**Supplementary Figure 4. Analysis of the SHOC2-PP1CA interface. a** The proposed SILK and RVxF binding motifs mapped onto SHOC2 (pink spheres) of the SMP complex. **b** Superposition of all RVxF-PP1C complexes present in the PDB onto the SMP complex. Surface of PP1CA (green) withthe RVxF motif of SHOC2 (current work, pink), muscle glycogen-targeting subunit (PDB ID 6DNO, cyan, Kumar, G.S. *et al*., (2018). Sci Adv. 4: eaau6044), RepoMan (PDB ID 5IOH, magenta, Kumar, G.S. *et al*., (2016). Elife 5: 16539 ), cell-permeable peptide (PDB ID 4G9J, salmon, Chatterjee, J. *et al*., (2012). Angew. Chem. Int. Ed. Engl. 51: 10054-10059), PP1 regulatory subunit 3A (PDB ID 5ZQV, light gray, Yu, J. *et al*., (2018). FEBS J. 285: 4646-4659), PP1 regulatory subunit 3B (PDB ID 5ZT0, violet, Yu, J. *et al*., (2018). FEBS J. 285: 4646-4659), Phactr1 (PDB ID 6ZEF, teal, Fedoryshchak, R.O. *et al.,* (2020). Elife 9: 61509), NIPP1 (PDB ID 3V4Y, orange, O'Connell, N. *et al*., (2012). Structure 20: 1746-1756), Retinoblastoma-associated protein (PDB ID 3N5U, purple, Hirschi, A. *et al*., (2010). Nat. Struct. Mol. Biol. 17: 1051-1057), PP1 regulatory subunit 10 (PDB ID 4MOY, gray, Choy, M.S. *et al*., (2014). Proc. Natl. Acad. Sci. 111: 4097-4102), GADD34 (PDB ID 4XPN, dark blue, Choy, M.S. *et al*., (2015). Cell. Rep. 11: 1885-1891), Spinophilin (PDB ID 3EGG, gold, Ragusa, M.J. *et al*., (2010). Nat. Struct. Mol. Biol. 17: 459-464) and mouse-inhibitor 2 (PDB ID 2O8G, dark olive, Hurley, T.D. *et al*., (2007) J. Biol. Chem. 282: 28874-28883). **c** Sequence alignment of the RVxF motif of SHOC2 across different species. Totally conserved residues are bold and highlighted in black, while similar residues are bold and highlighted in white. The RVxF motif is denoted with black stars. **d** Single-cycle kinetic analysis was performed on immobilized avi-tagged SHOC2 mutants as denoted in the figure with five injections of $MRAS_{GMPPNP}$ and PP1CA (blue). The data were fit to a 1:1 kinetic model (black) **e** The SILK binding pocket on PP1C (green surface), as shown by the SILK of mouse inhibitor-2 (yellow cartoon) is occluded by SHOC2 (pink). A hydrogen bond forms between E54 of PP1CA at the periphery of the SILK binding pocket to R203 of SHOC2.

a

b

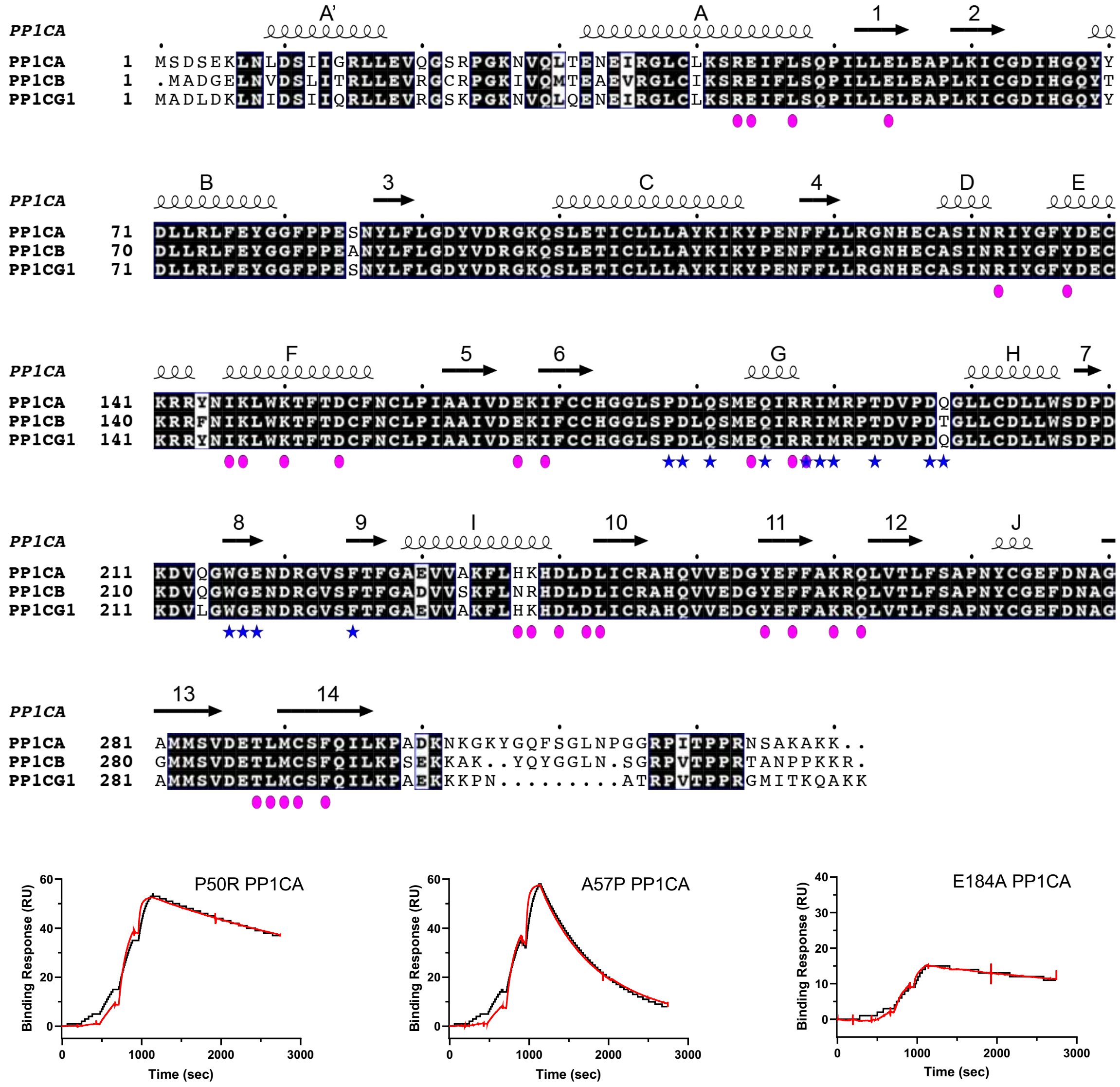

**Supplementary Figure 5. Isoforms and Noonan Syndrome mutations of PP1C. a** Sequence alignment of the three human isoforms of PP1C. Totally conserved residues are bold and highlighted in black, while similar residues are bold and highlighted in white. Non-conserved residues are only highlighted in white. The secondary structure of the PP1CA structure is shown above the alignment. α-helices and β-strands are labeled according to the nomenclature of Peti *et al.* (2012, *FEBS Journal* 280:596-611). Residues of PP1CA which interact with SHOC2 and MRAS are denoted with pink ovals and blue stars, respectively. R188 is the only residue of PP1CA which interacts with both SHOC2 and MRAS. **b** Single-cycle kinetic analysis was performed on immobilized avi-tagged SHOC2 with five injections of $MRAS_{GMPPNP}$ and PP1CA mutants as denoted in the figure (blue). The data were fit to a 1:1 kinetic model (black).

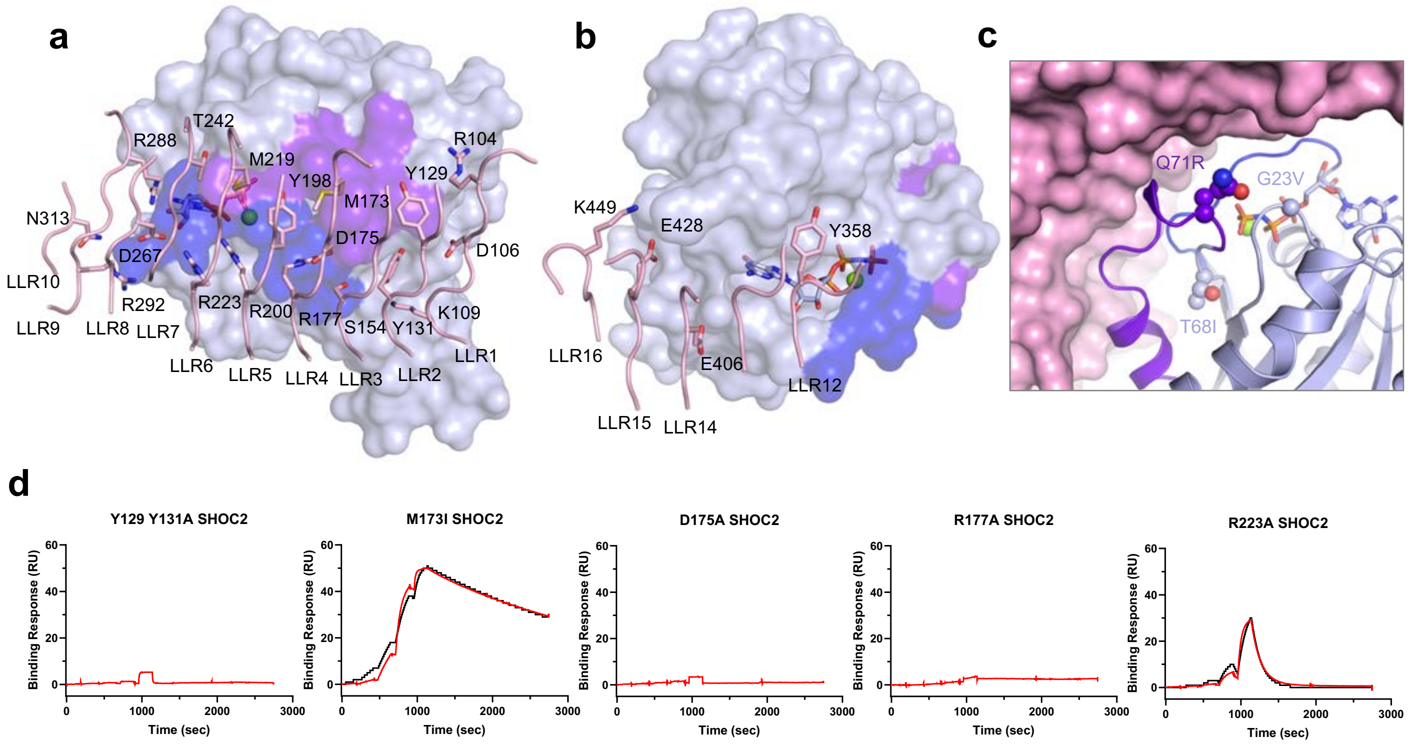

**Supplementary Figure 6. Analysis of SHOC2 binding to the surface of MRAS. a** The N-terminal LRRs of SHOC2 (pink) are shown interacting with the switch I (dark blue) and switch II (purple) of MRAS (blue). **b** The C-terminal LRRs of SHOC2 (pink) are shown interacting with the C-terminus of MRAS (blue surface). **c** Residues of MRAS found mutated in NS highlighted as spheres on the structure of MRAS. **d** Single-cycle kinetic analysis was performed on immobilized avi-tagged SHOC2 mutants as denoted in the figure with five injections of $MRAS_{GMPPNP}$ and PP1CA (blue). The data were fit to a 1:1 kinetic model (black).

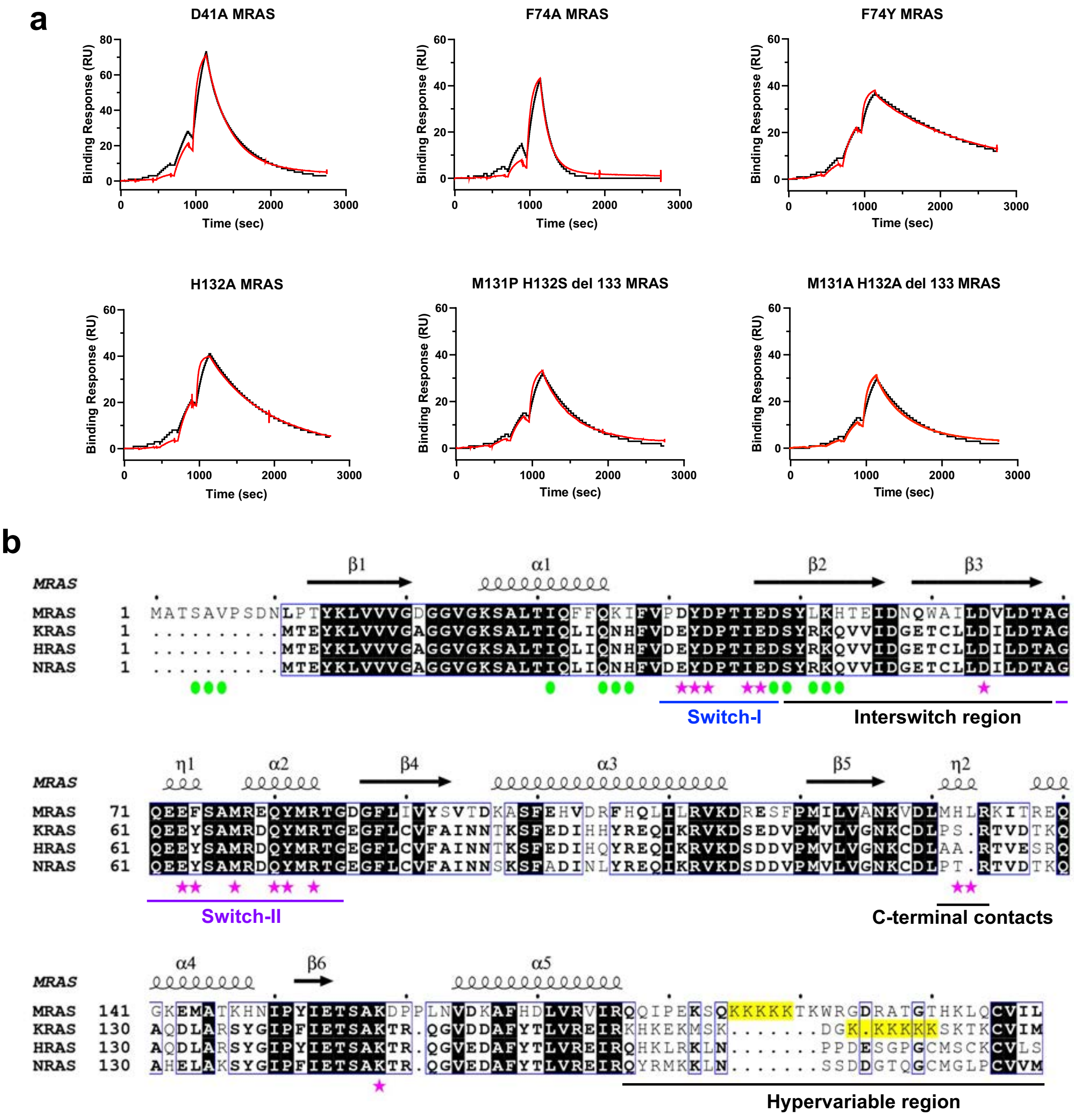

**Supplementary Figure 7 Analysis of MRAS binding to the surface of SHOC2. a** Single-cycle kinetic analysis was performed on immobilized avi-tagged SHOC2 with five injections of MRAS$_{\text{mutant-GMPPNP}}$ as denoted in the figure and PP1CA (blue). The data were fit to a 1:1 kinetic model (black). **b** Sequence alignment of human MRAS, KRAS, HRAS and NRAS sequences. Totally conserved residues are bold and highlighted in black, while highly conserved residues are bold and highlighted in white. Non-conserved residues are only highlighted in white. The secondary structure of the MRAS present in the SMP complex is shown above the alignment. Residues of MRAS which interact with SHOC2 and PP1CA are denoted with pink stars and green ovals, respectively.

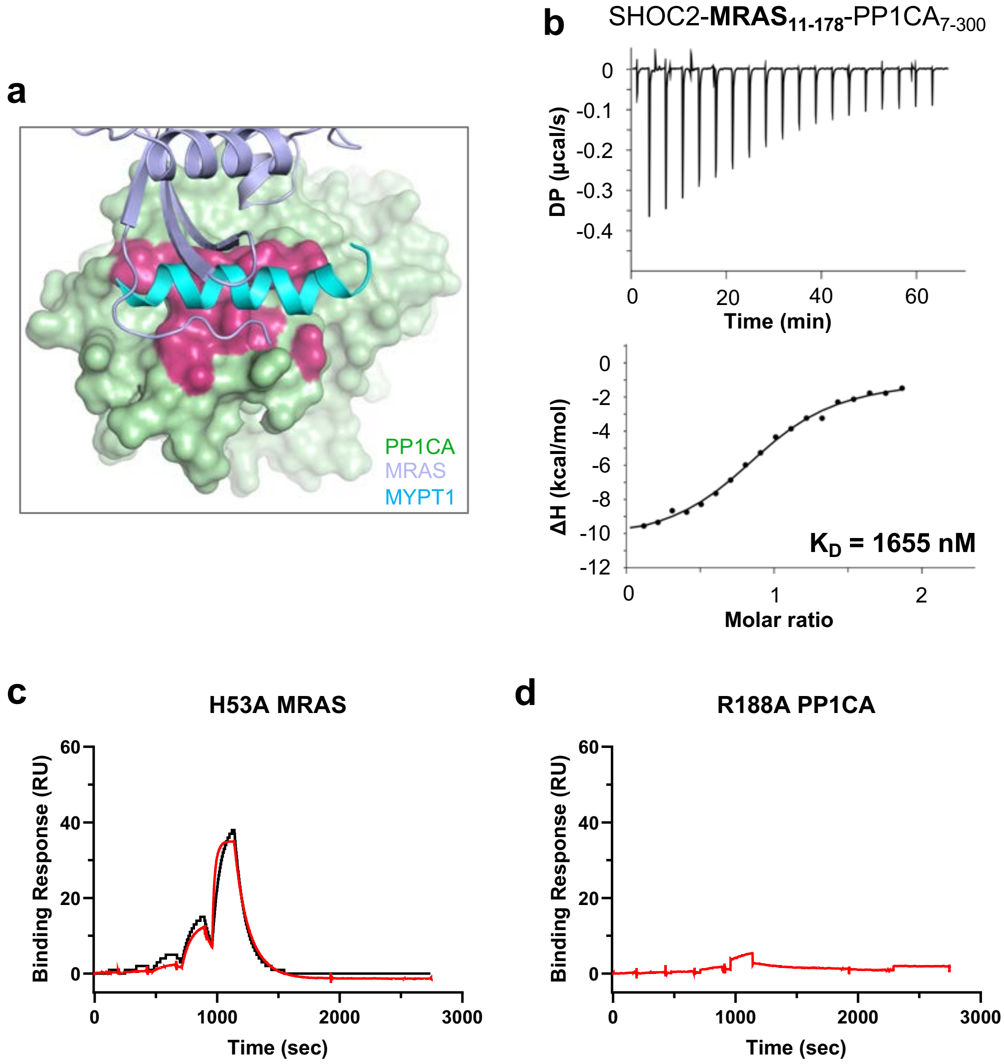

**Supplementary Figure 8. Analysis of the MRAS-PP1CA interface. a** The N-terminus of MRAS (blue) occupies the MyPhoNE cleft (dark red) on PP1CA (green). The helical MyPhoNE motif of MYPT1 (PDB ID 1s70) is shown in cyan. **b** ITC experiment to measure the dissociation constant between SHOC2, $MRAS_{11\text{-}178}$ and $PP1CA_{7\text{-}300}$ **c** Single-cycle kinetic analysis was performed on immobilized avi-tagged SHOC2 with five injections of $MRAS_{H53A\text{-}GMPPNP}$ and PP1CA (blue). **d** Single-cycle kinetic analysis was performed on immobilized avi-tagged SHOC2 with five injections of $MRAS_{GMPPNP}$ and $PP1CA_{R188A}$ (blue). In each case, the data were fit to a 1:1 kinetic model (black).

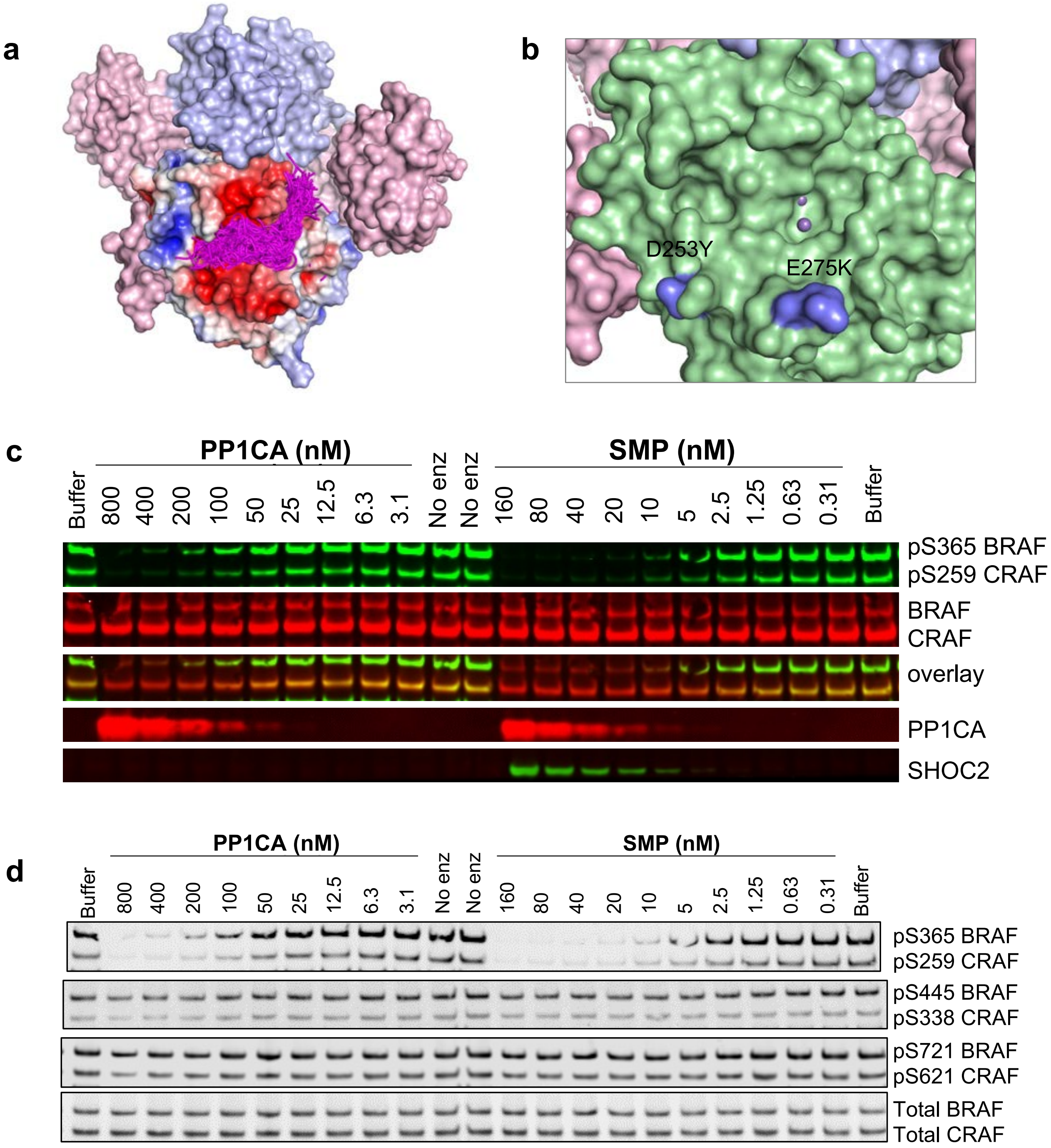

**Supplementary Figure 9. Docking of CRAF substrates and Noonan syndrome mutations found in the active site of PP1C. a** The CABS-dock server was used to generate a 15-mer peptide of the CR2-pS region of CRAF and dock into the PP1CA structure of the SMP complex. All 167 peptides were placed in the active site, with all peptides placed with the N- and C-termini in the acidic and hydrophobic active site channels (magenta ribbons). PP1CA is shown as an electrostatic surface. **b** Two NS mutations are found to line the acidic and C-terminal channels of PP1CB. These were mapped onto the PP1CA surface with D253Y and E275K shown in blue (D252Y and E274K in PP1CB). **c** Fluorescent Western blot of different concentrations of PP1CA or SMP incubated with either BRAF or CRAF substrates monitoring loss of CR2-pS signal (top gel, green bands). Total RAF loaded are shown as red bands (2nd from top gel). **d** Western blot of different concentrations of PP1CA or SMP with either BRAF or CRAF substrates. Blots were probed at different phosphorylation sites in the substrates. SMP complex only dephosphorylates CR2-pS.

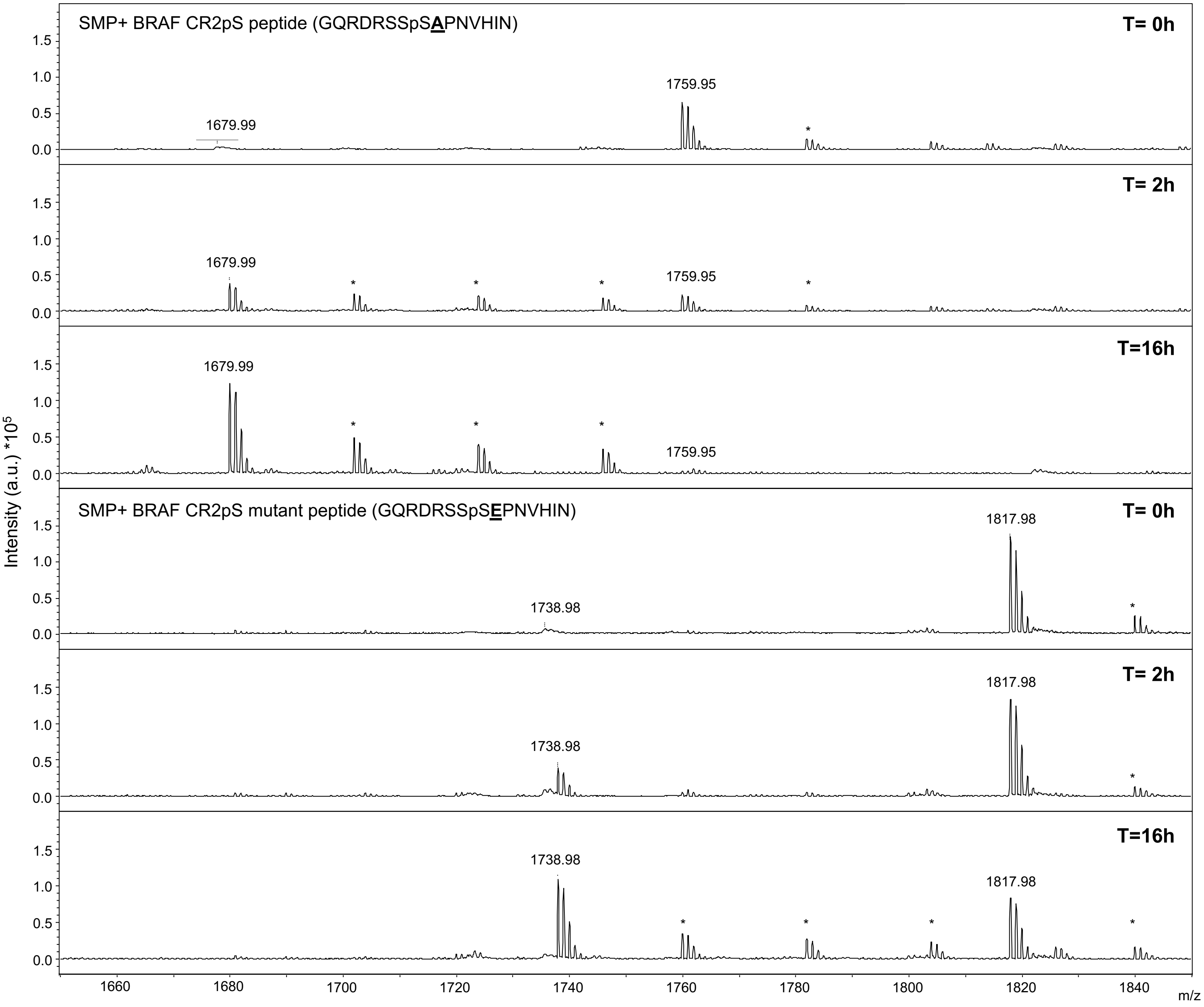

**Supplementary Figure 10. Dephosphorylation of BRAF CR2-pS phosphopeptides by the SMP complex.** Dephosphorylation of BRAF CR2-pS 15mer wild type peptide (top) and +1-position mutation to glutamic acid (bottom) as monitored by MALDI-TOF over 16 hours by the SMP complex. Sodium adducts of both dephosphorylated and phosphorylated peptides are denoted with *.

**Supplementary Table 1. Data collection and refinement statistics.**

| | SMP complex | SHOC2 |
|---|---|---|
| **Data collection** | | |
| Space group | P41 21 2 | P32 |
| Cell dimensions | | |
| *a*, *b*, *c* (Å) | 129.94, 129.94, 326.80 | 77.57, 77.57, 83.01 |
| $\alpha, \beta, \gamma$ (°) | 90.0, 90.0, 90.0 | 90.0, 90.0, 120.0 |
| Resolution (Å) | 163.40-2.17(2.20-2.17) * | 41.51-2.40(2.49-2.40) |
| $R_{merge}$ | 0.137(2.007) | 0.036(1.090) |
| $R_{pim}$ | 0.077(1.228) | 0.024(0.721) |
| $I / \sigma I$ | 8.5(0.9) | 19.8(1.7) |
| Completeness (%) | 99.9(98.4) | 99.9(100.0) |
| Redundancy | 7.9(6.8) | 6.2(6.4) |
| CC(1/2) | 0.997(0.340) | 1.000(0.581) |
| **Refinement** | | |
| Resolution (Å) | 101.7-2.17 | 35.31-2.40 |
| No. reflections | 148297 | 21811 |
| $R_{work}$ / $R_{free}$ | 19.6/22.6 | 21.7/26.7 |
| No. atoms | | |
| Protein | 15683 | 3742 |
| Ligand/ion | 180 | 31 |
| Water | 1046 | 36 |
| *B*-factors | | |
| Protein | 51.3 | 86.6 |
| Ligand/ion | 70.0 | 122.2 |
| Water | 52.8 | 86.0 |
| R.M.S. deviations | | |
| Bond lengths (Å) | 0.002 | 0.004 |
| Bond angles (°) | 0.51 | 0.69 |

* Values in the parenthesis are for the highest-resolution shell.

### Supplementary Table 2. SPR binding data for the formation of SHOC2-RAS-PP1CA complexes

| Immobilized ligand | Analyte | $k_{a\ mean}$ (1/Ms) | $k_a$ SD (1/Ms) | $k_{d\ mean}$(1/s) | $k_d$ sd (1/s) | $K_{D\ Mean}$ (M) | $K_D$ SD (M) | n |
|---|---|---|---|---|---|---|---|---|
| Avi SHOC2 | WT MRAS + WT PP1CA | 8.31E+03 | 6.85E+02 | 9.83E-04 | 1.40E-04 | 1.19E-07 | 1.68E-08 | 8 |
| Avi SHOC2 | Q71L MRAS + WT PP1CA | 1.57E+04 | | 4.10E-04 | | 2.60E-08 | | 1 |
| Avi SHOC2 | WT KRAS + WT PP1CA | 7.96E+0.4 | 1.04E+04 | 5.62E-02 | 7.87E-03 | 7.23E-07 | 9.63E-08 | 2 |
| Avi SHOC2* | WT HRAS + WT PP1CA | | | | | 1.52E-06 | 5.50E-07 | 2 |
| Avi SHOC2* | WT NRAS + WT PP1CA | | | | | 3.79E-06 | 1.58E-06 | 2 |
| **SHOC2 mutants** | | | | | | | | |
| Avi SHOC2 V64A | WT MRAS + WT PP1CA | 2.17E+02 | 1.32E+02 | 1.27E-02 | 1.91E-03 | 7.52E-05 | 5.46E-05 | 2 |
| Avi SHOC2 Y66A | WT MRAS + WT PP1CA | 2.82E+02 | 6.77E+01 | 2.04E-02 | 2.59E-03 | 7.50E-05 | 1.89E-05 | 3 |
| Avi SHOC2 Y129A/Y131A | WT MRAS + WT PP1CA | | | | | NB | | 3 |
| Avi SHOC2 E155A* | WT MRAS + WT PP1CA | | | | | 1.08E-06 | 6.63E-08 | 2 |
| Avi SHOC2 M173I | WT MRAS + WT PP1CA | 1.68E+04 | 8.02E+02 | 3.66E-04 | 1.18E-05 | 2.19E-08 | 1.09E-09 | 2 |
| Avi SHOC2 D175N | WT MRAS + WT PP1CA | | | | | NB | | 3 |
| Avi SHOC2 R177A | WT MRAS + WT PP1CA | | | | | NB | | 3 |
| Avi SHOC2 R223A | WT MRAS + WT PP1CA | 2.65E+02 | 5.35E+00 | 8.74E-03 | 1.14E-03 | 3.30E-05 | 4.96E-06 | 2 |
| **MRAS mutants** | | | | | | | | |
| Avi SHOC2 | D41A MRAS + WT PP1CA | 1.75E+03 | 2.65E+02 | 3.22E-03 | 9.15E-04 | 1.81E-06 | 2.47E-07 | 3 |
| Avi SHOC2 | H53A MRAS + WT PP1CA | 5.21E+03 | 1.12E+03 | 9.45E-03 | 1.70E-03 | 1.89E-06 | 7.33E-07 | 3 |
| Avi SHOC2 | F74A MRAS + WT PP1CA | 2.25E+02 | 3.77E+01 | 7.99E-03 | 1.03E-03 | 3.57E-05 | 1.42E-06 | 3 |
| Avi SHOC2 | F74Y MRAS + WT PP1CA | 9.06E+03 | 7.11E+02 | 7.02E-04 | 1.60E-05 | 7.78E-08 | 7.86E-09 | 2 |
| Avi SHOC2 | M131P H132S del 133 MRAS + WT PP1CA | 5.99E+03 | 3.56E+02 | 1.93E-03 | 4.84E-05 | 3.23E-07 | 2.72E-08 | 2 |
| Avi SHOC2 | M131A H132A del133 MRAS + WT PP1CA | 4.50E+03 | 1.43E+02 | 1.91E-03 | 1.46E-04 | 4.25E-07 | 1.89E-08 | 2 |
| Avi SHOC2 | H132A MRAS + WT PP1CA | 6.94E+03 | 0.00E+00 | 1.85E-03 | 7.73E-04 | 2.74E-07 | 1.00E-07 | 3 |
| **PP1CA mutants** | | | | | | | | |
| Avi SHOC2 | WT MRAS + P50A PP1CA | 8.68E+03 | 3.80E+03 | 2.52E-04 | 2.23E-05 | 3.27E-08 | 1.69E-08 | 2 |
| Avi SHOC2 | WT MRAS + A57P PP1CA | 8.73E+03 | 2.25E+03 | 1.17E-03 | 1.17E-04 | 1.37E-07 | 2.19E-08 | 2 |
| Avi SHOC3 | WT MRAS + E184A PP1CA | 5.67E+03 | 3.12E+03 | 1.75E-04 | 3.12E-05 | 3.45E-08 | 1.35E-08 | 2 |
| Avi SHOC2 | WT MRAS + R188A PP1CA | | | | | NB | | 2 |

n=number of replicates
*Steady-state analysis
NB – No binding

### Supplementary Table 3. ITC binding data for the formation of SHOC2-MRAS-PP1C complexes

| Syringe | Cell | $K_{D\ mean}$ (nM) | $K_D$ SD (nM) | ΔH (kcal $mol^{-1}$) | ΔH SD (kcal $mol^{-1}$) | -TΔS (kcal $mol^{-1}$) | -TΔS SD (kcal $mol^{-1}$) | ΔG (kcal $mol^{-1}$) | ΔG SD (kcal $mol^{-1}$) | n |
|---|---|---|---|---|---|---|---|---|---|---|
| SHOC2 | $MRAS_{1-178}$ + $PP1CA_{7-300}$ | 352 | 70 | -9.23 | 0.21 | 0.48 | 0.32 | -8.81 | 0.12 | 2 |
| SHOC2 | $MRAS_{1-178}$ + $PP1CA_{2-330}$ | 360 | 62 | -10.04 | 0.37 | 1.23 | 0.47 | -8.70 | 0.04 | 2 |
| SHOC2 | $MRAS_{1-178}$ + $PP1CB_{2-327}$ | 282 | 10 | -10.60 | 0.71 | 1.65 | 0.65 | -8.94 | 0.02 | 2 |
| SHOC2 | $MRAS_{11-178}$ + $PP1CA_{7-300}$ | 1655 | 262 | -9.45 | 0.54 | 1.55 | 0.64 | -7.89 | 0.10 | 2 |

n=number of replicates
NB – No binding
